# Differential antitumor activity of compounds targeting the ubiquitin-proteasome machinery in gastrointestinal stromal tumor (GIST) cells

**DOI:** 10.1101/791426

**Authors:** Jessica L. Rausch, Areej A. Ali, Donna M. Lee, Yemarshet K. Gebreyohannes, Keith R. Mehalek, Aya Agha, Sneha S. Patil, Yanis Tolstov, Jasmien Wellens, Harbir S. Dhillon, Kathleen R. Makielski, Maria Debiec-Rychter, Patrick Schöffski, Agnieszka Wozniak, Anette Duensing

**Author notes:** Corresponding author: Anette Duensing, MD; UPMC Hillman Cancer Center; Research Pavilion, Suite G.17; 5117 Centre Avenue; Pittsburgh, PA 15213; USA.

## Abstract

The majority of gastrointestinal stromal tumors (GISTs) are driven by oncogenic KIT signaling and can therefore be effectively treated with the tyrosine kinase inhibitor (TKI) imatinib mesylate. However, most GISTs develop imatinib resistance through secondary *KIT* mutations. The type of resistance mutation determines sensitivity to approved second-/third-line TKIs but shows high inter- and intratumoral heterogeneity. Therefore, therapeutic strategies that target KIT independently of the mutational status are intriguing. Inhibiting the ubiquitin-proteasome machinery with bortezomib is effective in GIST cells through a dual mechanism of *KIT* transcriptional downregulation and upregulation of the pro-apoptotic histone H2AX but clinically problematic due to the drug’s adverse effects. We therefore tested second-generation inhibitors of the 20S proteasome (delanzomib, carfilzomib and ixazomib) with better pharmacologic profiles as well as compounds targeting regulators of ubiquitination (b-AP15, MLN4924) for their effectiveness and mechanism of action in GIST. All three 20S proteasome inhibitors were highly effective *in vitro* and *in vivo*, including in imatinib-resistant models. In contrast, b-AP15 and MLN4924 were only effective at high concentrations or had mostly cytostatic effects, respectively. Our results confirm 20S proteasome inhibitors as promising strategy to overcome TKI resistance in GIST, while highlighting the complexity of the ubiquitin-proteasome machinery as a therapeutic target.

## INTRODUCTION

The majority of gastrointestinal stromal tumors (GISTs) are driven by a constitutively activating mutation in the *KIT* or *PDGFRA* (platelet-derived growth factor receptor alpha) receptor tyrosine kinase. While the tyrosine kinase inhibitor (TKI) imatinib mesylate (Gleevec®) is a highly effective first-line drug for inoperable or metastatic GIST, resistance occurs in approximately 50% of patients within the first two years of treatment^1^. Because the FDA-approved second- and third-line drugs sunitinib and regorafenib oftentimes offer little additional benefit, there is a need for new therapeutic approaches^2^. The major mechanism of TKI resistance involves secondary mutations in the primarily affected kinase indicating the continued dependency on KIT/PDGFRA activation. Therefore, therapeutic strategies targeting these kinases without the need of kinase domain binding seem particularly promising.

The 26S proteasome is a 2.5 MDa multiprotein complex and the main protein degradation machinery of eukaryotic cells^3^. It consists of a 20S tube-like proteolytic core particle and two 19S regulatory particles at either end. Proteins destined to be degraded are selectively targeted to the proteasome by the addition of a series of covalently attached ubiquitin molecules. Deubiquitinating enzymes (DUBs) associated with the 19S regulatory subunit remove these ubiquitin chains before proteins can enter the proteolytic subunit. The 20S core contains three major proteolytic activities (β5 chymotrypsin-like, β1 caspase-like, β2 trypsin-like). Inhibitors of the 20S proteasome core particle, such as the prototype proteasome inhibitor bortezomib (Velcade®), have gained clinical importance for the treatment of multiple myeloma and certain lymphomas^4^.

Previous studies from our laboratory have shown that targeting the ubiquitin-proteasome machinery with bortezomib is highly effective in GIST cells^5^. We could demonstrate that bortezomib-induced apoptosis is mediated by a dual mechanism of action: increased levels of soluble, non-chromatin-bound pro-apoptotic histone H2AX and a dramatic downregulation of KIT expression mediated by inhibition of active gene transcription^5–7^. It is known that loss of KIT expression is a strong inducer of apoptosis in GIST cells^6,8^. Although bortezomib has not shown significant clinical activity in many solid tumors, including an array of sarcomas^9^, there are recent reports of its clinical activity in GIST. For example, in a study evaluating bortezomib as a daily subcutaneous regimen in various solid tumors, the most significant response was seen in a patient with GIST^10^. In another study testing bortezomib in combination with vorinostat, one of the two GIST patients achieved stable disease (SD)^11^. Nevertheless, bortezomib is associated with marked adverse effects, most importantly irreversible neuropathy, as well as an intravenous route of administration warranting the evaluation of second-generation proteasome inhibitors in GIST^10,12^.

Carfilzomib (Kyprolis®, PR-171), ixazomib (Ninlaro®, MLN-9708), and delanzomib (CEP-18770) are inhibitors of the 20S proteolytic core particle of the 20S proteasome, like bortezomib^4^. Carfilzomib was approved by the FDA in 2012 for therapy-resistant multiple myeloma and inhibits the β5 chymotrypsin-like subunit of the proteasome, similar to bortezomib, but does so irreversibly and with a higher selectivity^13,14^. Carfilzomib has been shown to have less off-target effects as well as a lesser degree of adverse effects^13,15^. Ixazomib is the first orally bioavailable inhibitor of the 20S proteasome^16^. It is a structural derivative of bortezomib with improved pharmacologic properties and reversibly inhibits the chymotrypsin-like β5 subunit of the 20S proteasome^14–16^. Ixazomib was recently approved for the treatment of multiple myeloma^17^. Delanzomib reversibly binds the proteasome and can be administered orally and intravenously^18,19^. It potently inhibits the β5 chymotrypsin-like and the β1 caspase-like subunit and exhibits a more sustained inhibition of proteasome activity in multiple myeloma cells when compared to bortezomib^18,19^. Results of a phase I/II clinical trial in multiple myeloma were recently reported^20,21^.

In addition to inhibitors of the proteolytic 20S core of the proteasome, several compounds targeting regulators of the proteasomal degradation process have recently emerged. As mentioned above, DUBs catalyze the cleavage of ubiquitin from ubiquitin-conjugated proteins at the 19S regulatory core particle of the 26S proteasome^3^. The DUB inhibitor b-AP15 interferes with the activity of USP14 (ubiquitin-specific peptidase 14) and UCHL5 (ubiquitin carboxyl-terminal hydrolase isozyme L5)^21,22^ resulting in rapid accumulation of high molecular weight ubiquitin conjugates and a functional shutdown of the proteasome^22^. b-AP15 has shown pre-clinical activity in multiple myeloma and Waldenström’s macroglobulinaemia^23,24^.

A second important regulatory process of the proteolytic machinery involves the small ubiquitin-like modifier NEDD8^25^. NEDD8 is attached to lysine residues of proteins in a similar fashion as ubiquitin, and “neddylation” has been specifically implicated in regulating the turnover of cell cycle-regulatory proteins, such as p27 ^Kip1 26,27^. Inhibition of neddylation leads to reduced protein degradation *via* reduced ubiquitination and proteasome targeting, while proteasome activity remains intact. MLN4924 (Pevonedistat®), the first-in-class inhibitor of this process that targets the NEDD8 activating enzyme (NAE), has shown pre-clinical activity in various malignancies and is currently tested in several clinical trials^28^.

In the present study, we could show that all three inhibitors of the 20S proteolytic core of the proteasome tested were highly effective in *KIT*-mutant GIST cells, independently from mutational status or imatinib sensitivity. Mechanistically, the pro-apoptotic activity was mediated by a dual mechanism identical to bortezomib that resulted in upregulation of histone H2AX and transcriptional inhibition of *KIT*. Of note, delanzomib was the compound effective at the lowest dose with a profile similar to bortezomib. By contrast, the DUB inhibitor b-AP15 and the NAE inhibitor MLN4924 had a substantially lesser effect. In summary, targeting the proteolytic 20S core unit of the proteasome in GIST could be a promising therapeutic strategy in TKI-resistant GISTs.

## RESULTS

### Second-generation inhibitors of the 20S proteasome carfilzomib, ixazomib and delanzomib effectively induce cell cycle arrest and apoptosis in GIST cells

To determine whether carfilzomib, ixazomib and delanzomib have an effect on GIST cell viability and/or apoptosis, imatinib (IM)-sensitive (GIST882, *KIT* K642E; GIST-T1, *KIT* V560_Y578del) and IM-resistant (GIST48, primary *KIT* V560D/secondary *KIT* D820A; GIST430, primary *KIT* V560_L576del/secondary *KIT* V654A) GIST cell lines were treated with increasing concentrations covering a 100,000-fold concentration range (0.1 nM – 10.0 μM). Our study focused entirely on *KIT*-mutant GIST, because unfortunately no appropriate models for *PDGFRA*-mutant or *KIT/PDGFRA*-wildtype GIST exist to date. Cell viability and caspase 3/7 activity were measured using luminescence-based assays (Fig. 1A, B; Suppl. Methods). All three compounds were highly effective, with IC_50_s as low as 9 nM (Table 1A). Delanzomib was most potent (IC_50_s of 9-36 nM) with a profile similar to bortezomib, whereas carfilzomib and ixazomib were approximately one order of magnitude less potent (IC_50_s of 67-1679 nM and 41-175 nM, respectively; Table 1A). Notably, drug response was independent of *KIT* mutational status and sensitivity to imatinib. In fact, IM-resistant cells had an overall lower IC_50_ for all drugs than IM-sensitive cells.

**Table 1.** IC_50_ and EC_50_ values of compounds affecting the ubiquitin proteasome machinery in imatinib (IM)-sensitive and IM-resistant GIST cells. **(A)** IC_50_ (cell viability) of carfilzomib (CFZ), ixazomib (IXA) and delanzomib (DLZ) in comparison to bortezomib (BO). **(B)** EC_50_s of 20S inhibition of carfilzomib (CFZ), ixazomib (IXA) and delanzomib (DLZ) in comparison to bortezomib (BO). **(C)** IC_50_s (cell viability) of the USP14/UCHL5 inhibitor b-AP15 and the NEDD8 activating enzyme inhibitor MLN4924. IM-sensitive: GIST882, GIST-T1; IM-resistant: GIST48, GIST430.

|  | IC<sub>50</sub> (nM) |  |  |  |
| --- | --- | --- | --- | --- |
|  | CFZ | IXA | DLZ | BO |
| GIST882 | 184 | 82 | 23 | 13 |
| GIST-T1 | 1,679 | 175 | 36 | 26 |
| GIST48 | 67 | 41 | 9 | 7 |
| GIST430 | 185 | 56 | 12 | 13 |

**(B)**
|  | EC<sub>50</sub> (nM) |  |  |  |
| --- | --- | --- | --- | --- |
|  | CFZ | IXA | DLZ | BO |
| GIST882 | 96 | 84 | 13 | 8 |
| GIST-T1 | 41 | 50 | 5 | 9 |
| GIST48 | 52 | 25 | 7 | 6 |
| GIST430 | 83 | 42 | 5 | 4 |

(C)
|  | IC<sub>50</sub> (nM) |  |
| --- | --- | --- |
|  | b-AP15 | MLN4924 |
| GIST882 | >10,000 | >10,000 |
| GIST-T1 | >10,000 | >10,000 |
| GIST48 | 473 | 7,333 |
| GIST430 | 1,219 | >10,000 |

**Figure 1.**
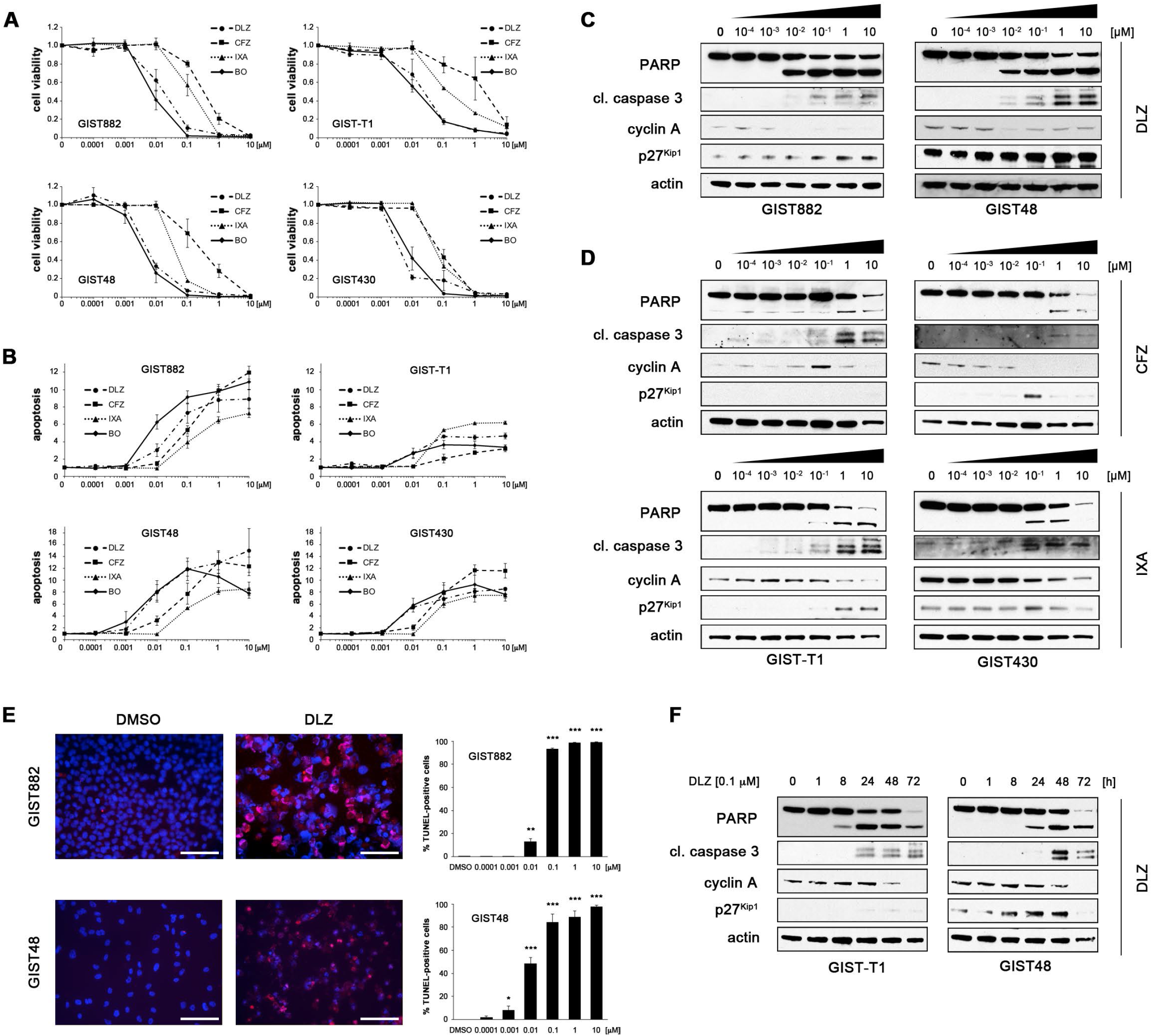
The second-generation inhibitors of the 20S proteasome delanzomib, carfilzomib and ixazomib effectively induce cell cycle arrest and apoptosis in imatinib (IM)-sensitive and IM-resistant GIST cells. **(A, B)** Dose-dependent effect of delanzomib (DLZ), carfilzomib (CFZ) and ixazomib (IXA) in comparison to bortezomib (BO) on cell viability **(A)** and induction of apoptosis **(B)** of imatinib (IM)-sensitive (GIST882, GIST-T1) and IM-resistant (GIST48, GIST430) GIST cells as measured by luminescence-based assays (mean +/-SE). **(C, D)** Immunoblot analysis for markers of cell cycle regulation and apoptosis after treatment of GIST cells with increasing concentrations (0.0001 μM – 10 μM; 72 h) of delanzomib **(C)**, carfilzomib **(D,** top panels) or ixazomib **(D,** bottom panels). **(E)** Dose-dependent effect of delanzomib on induction of apoptosis in GIST882 and GIST48 cells as measured by TUNEL assay. Cells were treated for 72 h. Graphs represent mean and standard error of at least three experiments with at least 100 cells counted each. Scale bars, 50 μm. *, p≤0.05 in comparison to control; **, p≤0.01 in comparison to control; ***, p≤0.001 in comparison to control (Student’s t-test, 2-tailed). **(F)** Immunoblot analysis for markers of cell cycle regulation and apoptosis after treatment of GIST cells with DMSO or delanzomib (0.1 μM) for the indicated times. **(C, D, F)** Grouped immunoblot images are either cropped from different parts of the same gel or from a separate gel run with another aliquot of the same protein lysate.

For the remainder of the manuscript, most *in vitro* assays were performed in all above-mentioned cell lines. For clarity and briefness, however, representative results of one IM-sensitive and one IM-resistant cell line are shown in the main figures, while all other data are included in the Supplemental Data section.

To biochemically assess tumor cell responses, we evaluated markers of apoptosis and cell cycle regulation by immunoblot analysis (Fig. 1C, D; Suppl. Fig. 1A; Suppl. Fig. 2A; Suppl. Methods). Delanzomib had a strong pro-apoptotic activity and also effectively induced cell cycle arrest at concentrations as low as 0.1 nM in IM-sensitive and IM-resistant cell lines (Fig. 1C, D; Suppl. Fig. 1A). In comparison, carfilzomib and ixazomib had a less pronounced effect (Fig. 1 D, Suppl. Fig. 2A), matching the results obtained in the luminescence-based assays (Fig. 1A, B).

The ability of the compounds to induce apoptosis was confirmed by the TUNEL assay and matched the immunoblotting studies (Fig. 1E, Suppl. Fig. 1B, Suppl. Fig. 2B, C; Suppl. Methods) with carfilzomib and ixazomib being less effective than delanzomib. Again, no general difference in activity between IM-sensitive and IM-resistant cells was noted. The effect of the compounds was time-dependent (Fig. 1F, Suppl. Fig. 1C, Suppl. Fig. 2D, E) with apoptosis starting to be evident as early as 8 h after treatment initiation (GIST-T1; Fig. 1F, Suppl. Fig. 2D, E). Interestingly, exit from the cell division cycle after delanzomib treatment lagged considerably. As before, no substantial difference between IM-sensitive and IM-resistant cells was seen.

Taken together, the second-generation 20S proteasome inhibitors delanzomib, carfilzomib and ixazomib showed significant cytotoxic activity in GIST cells, which was independent of the *KIT* mutational status and imatinib sensitivity. Delanzomib was the compound that was most effective at the lowest dose tested.

### The ability to inhibit the activity of the proteasome and induce accumulation of ubiquitinated proteins correlates with GIST cell cytotoxicity

Delanzomib, carfilzomib and ixazomib predominantly inhibit the chymotrypsin-like activity of the 20S proteasome in multiple myeloma cells^13,14,16,18,19,29,30^. To correlate 20S proteasome inhibitory activity with cytotoxicity in GIST cells, we measured the chymotrypsin-like activity of the 20S proteasome in a luminescence-based assay using Suc-LLVY-(succinyl-leucine-leucine-valine-tyrosine-)aminoluciferin as a substrate (Fig. 2A). While all three inhibitors strongly inhibited the proteasome, delanzomib was effective at the lowest dose, correlating with the compounds’ ability to induce apoptosis (EC_50_s of 5-13 nM, Table 1B). As before, no significant difference between IM-sensitive and IM-resistant GIST cells was seen. Interestingly, carfilzomib and ixazomib showed stronger proteasome inhibition capacity than delanzomib in the lower concentration range when treating IM-sensitive GIST882 and GIST-T1 cells. Together, our results indicate that the potency of proteasome inhibition indeed correlates with cytotoxic efficacy in GIST cells (Fig. 1; Suppl. Figs. 1, 2).

**Figure 2.**
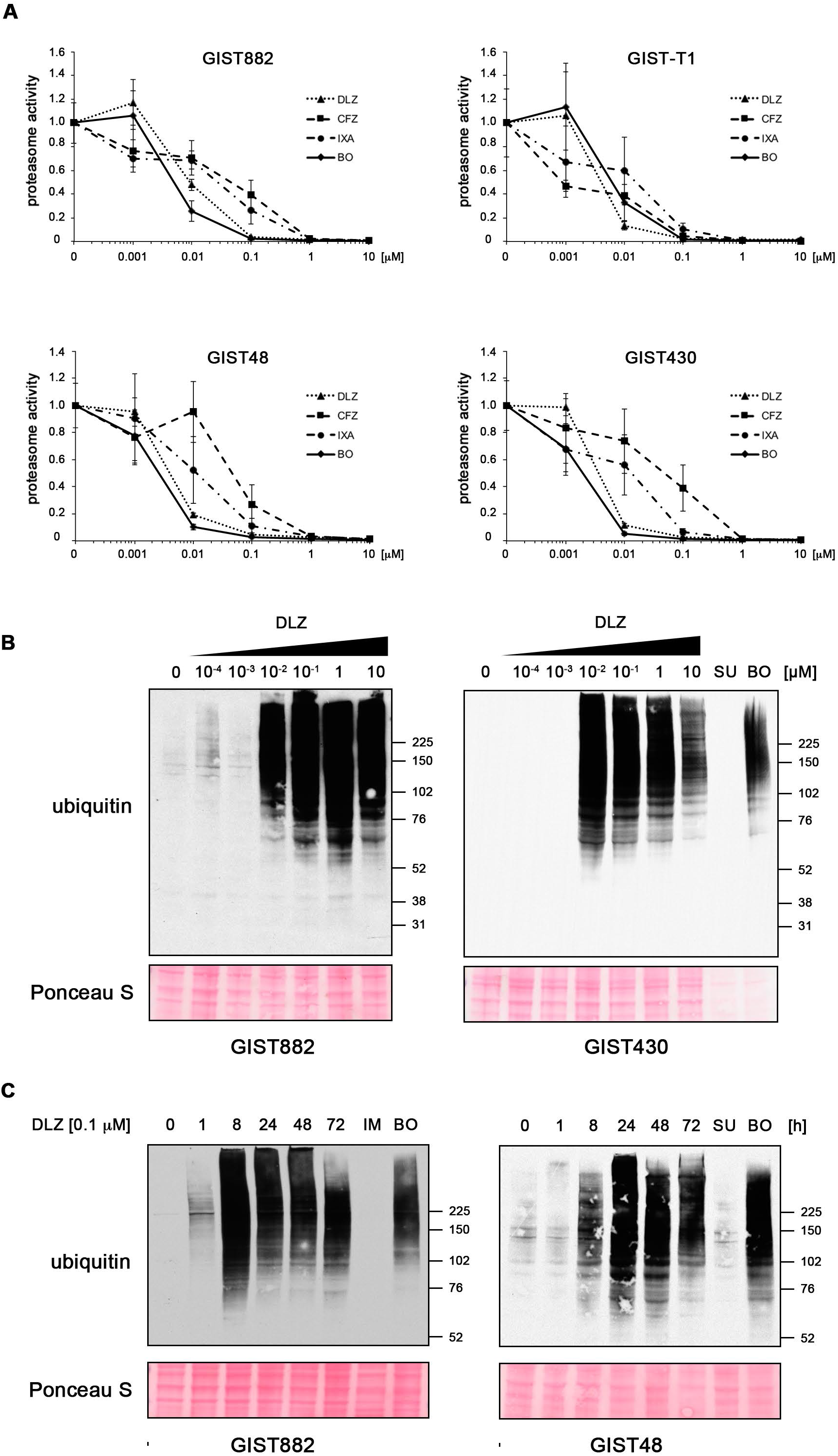
Delanzomib, carfilzomib and ixazomib strongly inhibit the activity of the 26S proteasome and rapidly lead to the accumulation of ubiquitinated proteins. **(A)** Dose-dependent effect of delanzomib (DLZ), carfilzomib (CFZ) and ixazomib (IXA) on the activity of the 26S proteasome in GIST cells. Cells were treated with increasing concentrations of inhibitor (0.0001 μM to 10 μM) for 2 h, and 26S proteasome activity was determined using a luminescence-based assay. **(B, C)** Dose- **(B)** and time-dependent **(C)** accumulation of mono-ubiquitinated proteins in GIST cells after delanzomib treatment as determined by immunoblotting. As expected, kinase inhibitor treatment with imatinib (IM) or sunitinib (SU; both 1.0 μM; 72 h) did not lead to increased ubiquitination. IM and SU serve as standard treatment controls for IM-naïve and IM-resistant cell lines, respectively. Bortezomib (BO) serves as control for proteasome inhibition.

In a next step, we assessed whether the accumulation of ubiquitinated proteins coincides with cytotoxicity of the compound. As expected, treatment with delanzomib led to a dose-dependent accumulation of ubiquitinated proteins and also had the strongest effect when compared to carfilzomib and ixazomib (Fig. 2B; Suppl. Fig. 1D, Suppl. Fig. 3A, C). This process was remarkably fast, starting after only one hour of treatment in most cell lines (Fig. 2 C; Suppl. Fig. 1E; Suppl. Fig. 3B, D). Kinase inhibitor treatments (imatinib in IM-sensitive cells, sunitinib in IM-resistant cells) that were used as controls did not lead to increased ubiquitination, as expected.

Taken together, delanzomib, carfilzomib and ixazomib effectively inhibit the chymotrypsin-like activity of the 20S proteasome in IM-sensitive and IM-resistant GIST cells and lead to the accumulation of ubiquitinated proteins. Delanzomib had the strongest effect of the three, suggesting that the cytotoxic effect of the compounds is indeed associated with their ability to inhibit the proteasome.

### Inhibition of the USP14/UCHL5 deubiquitinating enzymes or the NEDD8 activating enzyme (NAE) have a lower pro-apoptotic activity in GIST cells when compared to 20S inhibitors

Having shown that delanzomib, carfilzomib and ixazomib have activity against GIST cells, we tested whether inhibition of protein degradation by targeting regulators of the ubiquitin-proteasome pathway has the same effect. b-AP15 is a DUB inhibitor that interferes with ubiquitin cleavage at the 19S regulatory particle of the proteasome leading to a functional shutdown of the proteasome^3,20^. Significant reduction in cell viability and increase of apoptosis (luminescence-based assays) was only achieved after treatment with very high concentrations of b-AP15 (1 μM and higher; Fig. 3A). IC_50_s ranged from 0.5 μM and 1.2 μM in IM-resistant cells to over 10 μM in IM-sensitive cells (Table 1C). Immunoblot analysis paralleled these results (Fig. 3B; Suppl. Fig. 4A). While b-AP15 did lead to the accumulation of ubiquitinated proteins, concentrations of 1.0 μM or higher were needed to achieve this effect (Fig. 3C).

**Figure 3.**
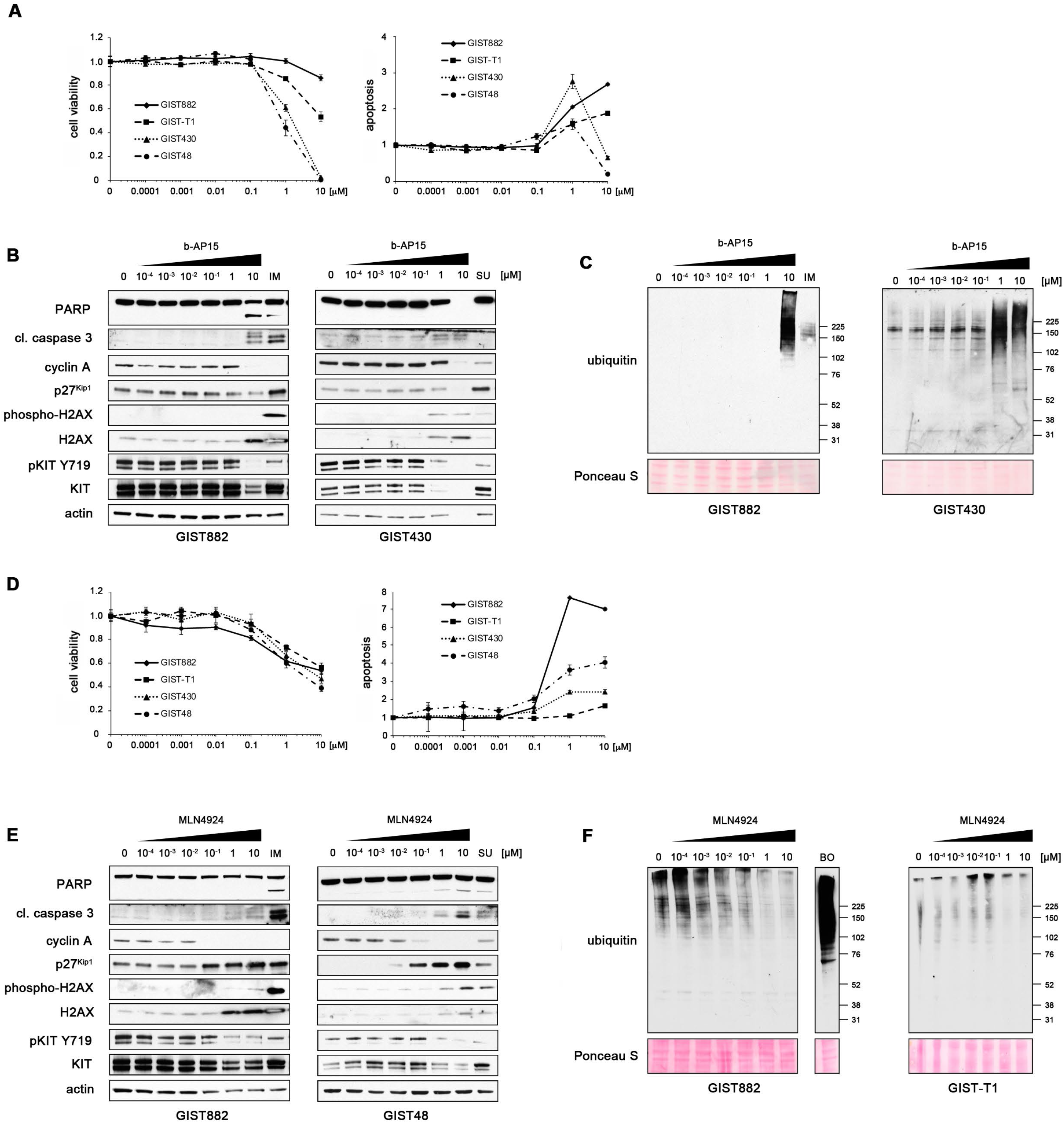
The USP14/UCHL5 inhibitor b-AP15 and the NAE inhibitor MLN4924 are only effective at high concentrations or have a mostly cytostatic effect in GIST cells. **(A)** Dose-dependent effect of b-AP15 on cell viability (left) and induction of apoptosis (right) of IM-sensitive (GIST882, GIST-T1) and IM-resistant (GIST48, GIST430) GIST cells as measured by luminescence-based assays (mean +/-SE). The abrupt drop of cells undergoing apoptosis in IM-resistant cells treated with 10 μM b-AP15 most likely indicates that the majority of cells were dead at this concentration and no caspase 3/7 activity (luminescence readout) was present. **(B, C)** Immunoblot analysis for markers of cell cycle regulation and apoptosis **(B)** and accumulation of mono-ubiquitinated proteins **(C)** in GIST cells after treatment with increasing concentrations (0.0001 μM – 10 μM; 72 h) of b-AP15. Treatment with IM or SU (both 1.0 μM) serve as standard treatment controls for IM-naïve and IM-resistant cell lines, respectively. **(D)** Dose-dependent effect of MLN4924 on cell viability (left) and induction of apoptosis (right) of GIST cells as measured by luminescence-based assays (mean +/-SE). **(E, F)** Immunoblot analysis for markers of cell cycle regulation and apoptosis **(E)** and accumulation of mono-ubiquitinated proteins **(F)** after treatment with increasing concentrations (0.0001 μM – 10 μM; 72 h) of MLN4924. Treatment with IM, SU (both 1.0 μM) or serve as standard treatment controls for IM-naïve and IM-resistant cell lines, respectively. Bortezomib (BO; 0.1 μM) serves as control for proteasome inhibition. **(B, E)** Grouped immunoblot images are either cropped from different parts of the same gel or from a separate gel run with another aliquot of the same protein lysate.

We next tested the NAE inhibitor MLN4924 for its effect on GIST cells. Neddylation is critical for the recruitment of the E2 ubiquitin-conjugating enzyme to the E3 ligase complex, and its inhibition decreases protein degradation by reducing ubiquitination, while leaving proteasome function intact^25,27^. Treatment of GIST cells with increasing concentrations of MLN4924 showed reduction of cellular proliferation starting at 0.1 μM but did not lead to a substantial apoptotic response (Fig. 3D). IC_50_s ranged from 7.3 μM to 461 μM (Table 1C), favoring IM-resistant cell lines. Results were paralleled on the molecular level by immunoblot analysis (Fig. 3E, Suppl. Fig. 4B). As expected from its mechanism of action, the amount of poly-ubiquitinated proteins decreased with increasing concentrations of MLN4924 (Fig. 3F). This effect started at fairly low concentrations (0.01 μM) and generally mirrored the cell cycle inhibitory action of the drug.

Taken together, both the DUB inhibitor b-AP15 and the NAE inhibitor MLN4924 only led to minimal pro-apoptotic responses in GIST cells, but MLN4924 showed a prominent cytostatic effect. Interestingly, b-AP15’s ability to induce apoptosis and MLN4924’s cell cycle inhibitory activity correlated with their ability to modulate ubiquitination and presumably inhibit protein degradation.

### Delanzomib, carfilzomib and ixazomib induce GIST cell death by upregulation of H2AX and transcriptional downregulation of KIT

As shown in an earlier study by our group, the prototype proteasome inhibitor bortezomib has a dual mechanism of action in GIST through upregulation of the pro-apoptotic histone H2AX and inhibition of transcription resulting in loss of KIT expression^5^. When analyzing H2AX levels after treatment with delanzomib, carfilzomib and ixazomib, we found that both phosphorylated and total histone H2AX protein expression increased in a dose- and time-dependent manner (Fig. 4A, B; Suppl. Fig. 5A, B; Suppl. Fig. 6A-D). This increase paralleled the accumulation of ubiquitinated proteins (Fig. 2B, C; Suppl. Fig. 1D, E; Suppl. Fig. 3) and was present in IM-sensitive and IM-resistant GIST cells. At the same time, KIT protein expression levels were substantially downregulated, and – as expected – KIT phosphorylation decreased in parallel (Fig.4A, B; Suppl. Fig. 5A, B; Suppl. Fig. 6A-D).

**Figure 4.**
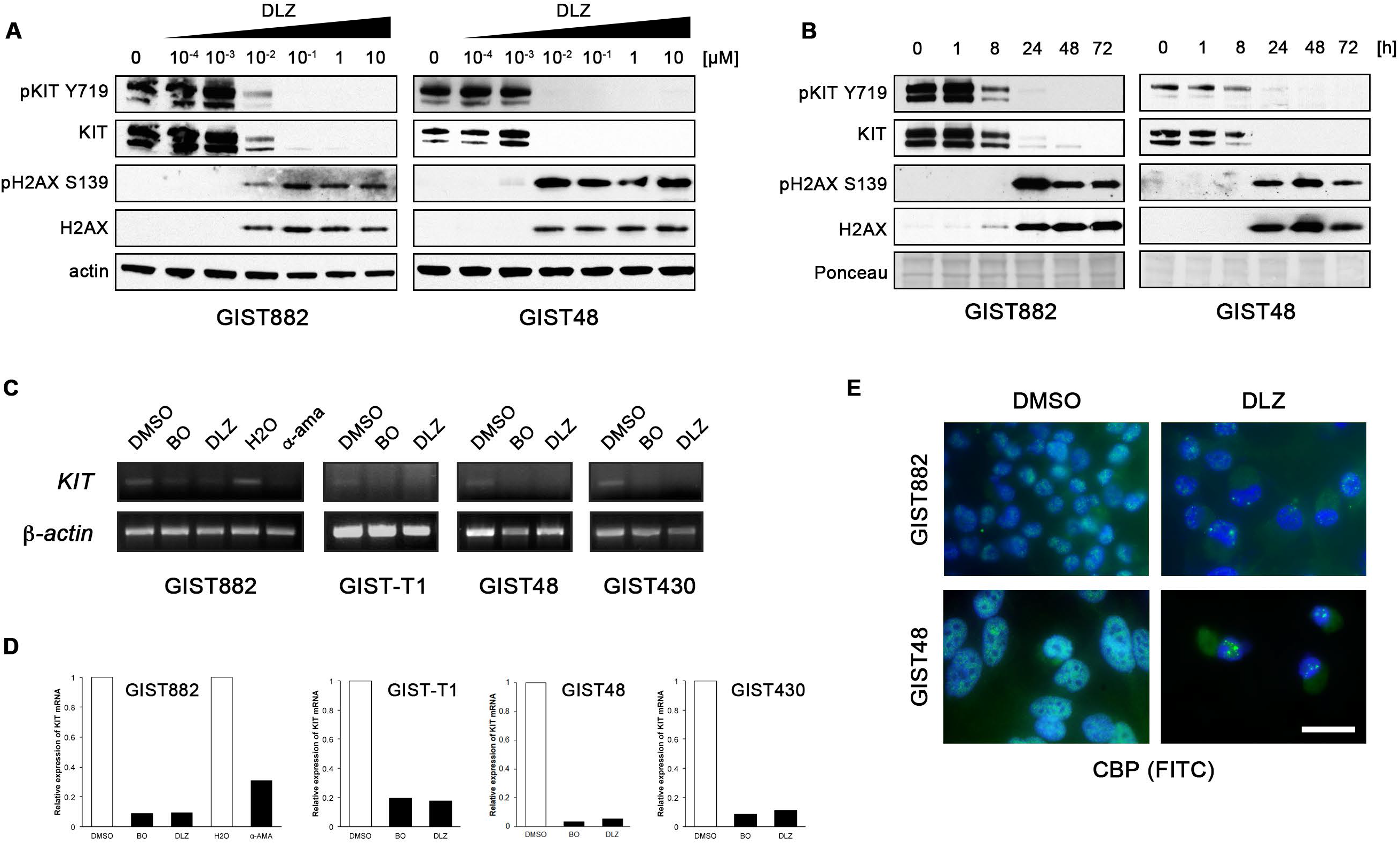
Delanzomib (DLZ) treatment leads to upregulation of soluble histone H2AX but downregulates KIT expression through inhibition of transcription. **(A, B)** Immunoblot analysis of GIST cells treated with delanzomib at the indicated concentrations for 72 h **(A)** or with 0.1 μM delanzomib for the indicated times **(B)** and probed for phospho-H2AX (S139) and total H2AX as well as phospho-KIT (Y719) and total KIT. Grouped immunoblot images are either cropped from different parts of the same gel or from a separate gel run with another aliquot of the same protein lysate. **(C, D)** RT-PCR **(C)** and quantitative RT-PCR (qRT-PCR) amplification **(D)** of *KIT* mRNA after treating GIST cells with DMSO or 0.1 μM delanzomib in comparison to bortezomib (BO) for 48 h. GIST882 cells were also treated with the RNA polymerase II inhibitor α-amanitin (α -ama) or H_2_O solvent control. **(E)** Immunofluorescence microscopic analysis of GIST882 and GIST48 cells treated with DMSO or 0.1 μM delanzomib for 72 h and stained for the transcriptional co-activator CREB-binding protein (CBP; green). Nuclei were stained with DAPI. Bar, 20 μm.

We next analyzed whether delanzomib leads to transcriptional inhibition of *KIT.* Indeed, *KIT* mRNA levels were substantially reduced as shown by RT-PCR (Fig. 4C; Suppl. Methods) and quantitative RT-PCR (qRT-PCR; Fig. 4D; Suppl. Methods). Downregulation of *KIT* transcripts was comparable to treatment with bortezomib or the highly effective RNA polymerase II inhibitor α-amanitin (Fig. 4C, D; left panels, GIST882).

Staining for the transcriptional co-activator CREB-binding protein (CBP) by immunofluorescence microscopy indicated that delanzomib affects GIST cell transcription in more general terms (Fig. 4E; Suppl. Fig. 5C; Suppl. Methods). In control-treated cells with active transcription, CBP localized to the transcriptional start sites resulting in a fine-speckled nuclear staining pattern but was redistributed into large nuclear displacement foci or was lost completely after delanzomib treatment, pointing towards a global transcriptional downregulation. Carfilzomib and ixazomib had a similar effect (Suppl. Fig. 6E, F). Transcriptional inhibition after delanzomib treatment was time-dependent over the course of 72 hours (Suppl. Fig. 7), correlating well with our immunoblotting data shown in Fig. 4. While the global transcriptional downregulation could be worrisome for non-specific toxicity, the fact that carfilzomib and ixazomib as well as bortezomib are safely used in the clinic, speaks against a major impact in patients.

Interestingly, treatment with b-AP15 and MLN4924 also led to increased phosphorylated/total H2AX levels as well as a reduction of phosphorylated/total KIT, however, only at the high drug concentrations that led to an apoptotic response (Suppl. Fig. 4). This is especially striking for MLN4924, because its ability to modulate ubiquitination levels and induce cell cycle exit requires much lower drug concentrations.

Taken together, the pro-apoptotic effect of all compounds tested in our study includes a dual mechanism of action with increased levels of histone H2AX and global transcriptional inhibition leading to loss of KIT expression. Nevertheless, there are vast differences in their efficacy.

### *Delanzomib has* in vivo *antitumor activity in GIST xenografts*

Since delanzomib was the strongest inducer of GIST cell death, we tested its *in vivo* antitumoral activity in several GIST xenograft models, including two patient-derived xenografts. Oral treatment with delanzomib led to significant responses in all models tested.

Because of the rapid kinetics seen in our *in vitro* studies, we performed a one-dose bolus experiment, in which mice bearing IM-resistant UZLX-GIST9 patient-derived xenografts (primary *KIT* P577del/secondary *KIT* W557LfsX5 and *KIT* D820G) received a single dose of oral delanzomib and were sacrificed 24 h later (Suppl. Fig. 8A). While no reduction in tumor size was seen or expected, histopathologic changes were readily detectable. This is especially important in light of the fact that delanzomib has been shown to rapidly distribute into the extra-vascular space after reaching peak concentration^31^. All treated tumors had a grade 2 or higher histologic response. These changes were accompanied by a significant decrease in the percentage of Ki-67-positive cells as well as a significant increase of cleaved PARP-positive cells by immunohistochemistry (Suppl. Methods). Some tumors already exhibited large areas of central necrosis.

Delanzomib treatment over the course of three weeks resulted in significant tumor reduction comparable to imatinib treatment in the IM-sensitive patient-derived UZLX-GIST1 model (*KIT* V560D) (Fig. 5A; Suppl. Fig. 8B)^32^. Tumor reduction after delanzomib was accompanied by a marked histopathologic response, and more than 60% of the responses were of grade 3 or higher (Fig. 5B). Response to delanzomib most prominently resulted in large areas of central necrosis, whereas control imatinib treatment rather led to myxoid degeneration, similar to what is seen in clinical imatinib-treated tumor samples^33^. Furthermore, delanzomib led to a highly significant reduction of proliferation, which was equivalent to imatinib treatment (Fig. 5B; reduction of the Ki-67-positive proliferation fraction from 26.8 % in placebo-treated controls to 3.0 % in delanzomib-treated tumors; decrease of phosphorylated histone H3 S10 positive mitotic cells from 88/10 HPF to 4/10 HPF). Delanzomib also resulted in increased apoptosis (cleaved PARP; Fig. 5B). While central areas of the tumor show massive cell death, the percentage of cleaved PARP-positive intratumoral cells (only intact cells counted) also increased (from 8.9 % in vehicle-treated controls to 15.0 % in delanzomib-treated animals), although not reaching statistical significance. Of note, no increase, but rather a reduced number of cleaved PARP-positive cells was detected in imatinib-treated tumors (1.6 %), indicating that the majority of the remaining tumor cells, while likely not proliferating, could be quiescent^34,35^.

**Figure 5.**
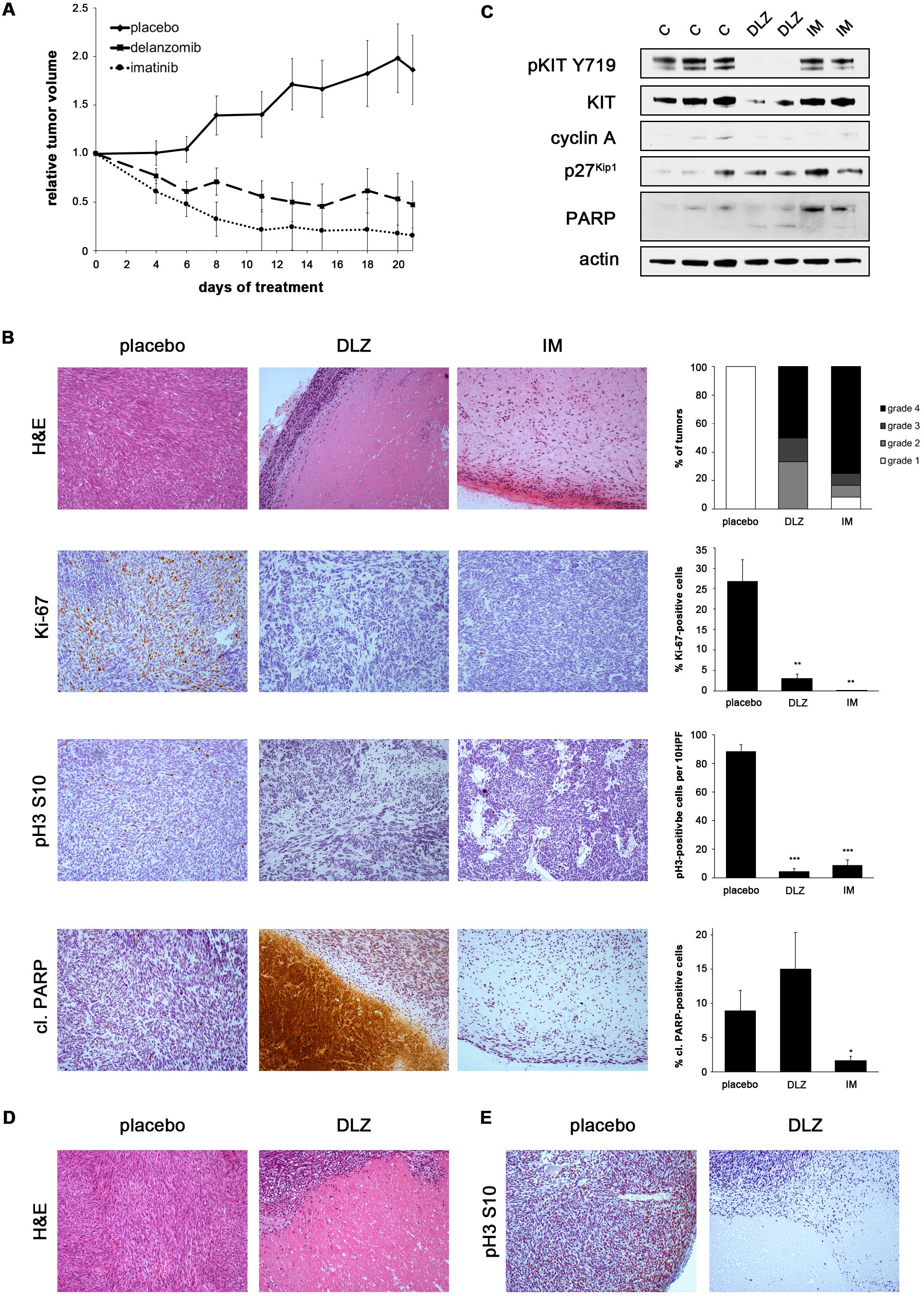
Delanzomib has *in vivo* antitumor activity in IM-sensitive and IM-resistant GIST xenografts. **(A)** Relative tumor volume of UZLX-GIST1 xenografts over the course of a 21-day treatment with vehicle, delanzomib or imatinib. Measurements represent the average of at least four tumors per group. **(B)** Histopathologic response of imatinib-sensitive UZLX-GIST1 xenografts to treatment with delanzomib in comparison with placebo-treated and imatinib-treated controls. Hematoxylin and eosin (H&E) as well as immunohistochemical staining for Ki-67, phosphorylated histone H3 S10 or cleaved PARP (10X magnification). Histopathologic response grade was determined according to Agaram et al.^55^. Data shown in graph represent at least six tumors per group. Quantification of proliferation index (% Ki-67 positive cells), mitotic cells (pH3-positive cells per 10 high-power fields, HPF) and percentage of cleaved PARP-positive cells (brown staining, respectively) represents the average of at least six tumors per group. It is of note, that only intact, individual cells were included in the counts. Hence, the central debris area in the cleaved PARP stain (although massively positive) is not reflected in the accompanying graph. Columns, mean + SE; *, p≤0.05 in comparison to control; **, p≤0.01 in comparison to control; ***, p≤0.001 in comparison to control (Student’s t-test, 2-tailed). **(C)** Immunoblot analysis of UZLX-GIST1 xenografts treated with delanzomib (24 h) in comparison with placebo and imatinib-treated positive control. Each lane represents one xenograft (C, vehicle control; DLZ, delanzomib; IM, imatinib). Grouped immunoblot images are either cropped from different parts of the same gel or from a separate gel run with another aliquot of the same protein lysate. **(D)** Histopathologic response of imatinib-resistant GIST430 xenografts to treatment with delanzomib in comparison with placebo. **(E)** Immunohistochemical staining for phosphorylated histone H3 S10 in GIST430 xenografts treated with delanzomib or placebo control.

The above changes were also reflected when assessing the response of UZLX-GIST1 to delanzomib (24 h) by immunoblotting (Fig. 5C). Interestingly, only the delanzomib-treated xenografts showed detectable PARP cleavage, while both delanzomib- and imatinib-treated xenografts exhibited reduced cyclin A levels and increased expression of p27^Kip1^ indicating cell cycle arrest. Importantly, delanzomib-treated xenografts also show a downregulation of total KIT protein expression, as would be expected from our *in vitro* data (Fig. 4). Interestingly, a moderate downregulation of KIT resulted in a complete loss of KIT activation. Results were very much comparable to the UZLX-GIST9 bolus treatment (Suppl. Fig. 8A) as both models already showed drastic changes after a single dose of delanzomib and a short follow-up.

We also tested delanzomib over the course of three weeks in IM-resistant GIST430 xenografts. While the reduction of tumor growth was not as dramatic as seen in the UZLX-GIST1 model (Suppl. Fig. 8C), the treatment nevertheless led to significant histopathologic changes (Fig. 5D) with 60% of the tumors showing a histopathologic response grade of 2 or higher (Suppl. Fig. 8D). Furthermore, treatment caused a highly significant reduction of proliferation (decrease of pH3-positive cells from 94/10 HPF 22.5/10 HPF; Fig. 5E, Suppl. Fig. 8E).

Taken together, delanzomib has significant activity in GIST xenografts, including IM-resistant models.

## DISCUSSION

Although most *KIT*-mutant GISTs can be successfully treated with tyrosine kinase inhibitors (TKIs), the majority of tumors develop therapy resistance over time, mostly through secondary *KIT* mutations. Those mutations show significant intertumoral and intratumoral heterogeneity and thereby confer sensitivity to different second- or third-line drugs. Therapeutically targeting KIT in a mutation-independent manner is hence an intriguing strategy.

In the present study, we show that targeting the proteolytic machinery is highly effective in *KIT*-mutant GIST cells and that activity of the tested compounds is independent of mutational status or sensitivity to imatinib. Apoptosis is induced through upregulation of the pro-apoptotic histone H2AX as well as transcriptional downregulation of *KIT*, a mechanism previously described by our group for the first-in-class proteasome inhibitor bortezomib^5–7^. Notably, all tested compounds share this mechanism, regardless of whether they target the 20S proteolytic core of the proteasome or other regulatory components. Of note, to our knowledge, there are no reports on the induction of H2AX levels by proteasome inhibitors in other disease entities, such as multiple myeloma.

The shared transcriptional downregulation of *KIT* in all compounds raises the possibility that it could be the result of increased H2AX levels, especially in light of the fact that non-chromatin-bound H2AX can lead to transcriptional inhibition^6^. However, we have shown previously that siRNA-mediated knockdown of H2AX did not completely rescue bortezomib-induced apoptosis, indicating that additional mechanisms are involved^6^. It is well-known that the proteasome has an important role in transcription initiation^36,37^. Transcriptional activation domains of many transcription factors, including ETV1, a main transcription factor of KIT in the GIST/interstitial cell of Cajal lineage^38^, often overlap with degrons^36^, indicating that their degradation is crucial for transcriptional activation. The fact that it has been shown that bortezomib also leads to transcriptional downregulation of Sp1 (another transcription factor involved in the regulation of *KIT*^39^) as well as Notch in multiple myeloma and acute lymphocytic leukemia strengthens our hypothesis^40,41^.

Although all compounds showed some effect in GIST cells, vast differences in drug efficacy were seen, ranging over three orders of magnitude. Compounds that inhibit the 20S catalytic core of the proteasome (delanzomib, carfilzomib and ixazomib), like bortezomib, stood out as most potent. Importantly, these second-generation 20S inhibitors have better pharmacologic properties than bortezomib, including an oral route of administration for some, as well as less toxicity, which warrant their pre-clinical examination without formally determining the clinical utility of bortezomib in GIST^15,16,19,30^. Nevertheless, not all 20S inhibitors were equally effective in GIST cells. In fact, the activity of delanzomib was one order of magnitude higher than that of carfilzomib and ixazomib and largely identical to that of bortezomib. The reason for the superior efficacy of delanzomib does not seem to lie in a structural similarity to bortezomib (ixazomib, and not delanzomib, is a direct derivative of bortezomib^16,30^) or its proteasome binding dynamics (carfilzomib, but not delanzomib, irreversibly inhibits the proteasome^13–16,18,19^). It has recently become clear, however, that co-inhibiting the caspase-like subunit of the proteasome in addition to the chymotrypsin-like subunit enhances the antineoplastic activity of proteasome inhibitors^42^. Interestingly, both delanzomib and bortezomib potently inhibit the β1 caspase-like subunit, while carfilzomib and ixazomib do not^15,37^, indicating that this ability could be the reason for the superior activity of this compound.

Based on our preclinical results, delanzomib holds most promise to be effective in the clinical setting for GIST patients. Two early phase clinical trials in multiple myeloma and solid tumors reported that delanzomib treatment was feasible^20,31^. Importantly, no neurotoxicity, the most severe adverse effect of bortezomib, was reported. While a few partial responses were seen after delanzomib, however, the best result for most patients was stable disease^20,31^. Due to this unexpectedly low efficacy, further development of delanzomib for multiple myeloma was discontinued, and the future of delanzomib as a treatment for GIST is unclear^20^. Nevertheless, testing other 20S proteasome inhibitors could be worthwhile. One possibility is ixazomib, which has recently been FDA-approved and was the second-most effective drug in our study. Importantly, ixazomib is orally administered and would therefore be well-accepted in a patient population that is not otherwise acquainted with intravenously administered medications^16^. In addition, third-generation compounds have become available since we began our study. Especially marizomib (salinosporamide A; Triphase/Celgene), could be of interest because of its ability to inhibit the β1 caspase-like and β2 trypsin-like subunit of the proteasome in addition to the β5 chymotrypsin-like subunit^43^. Marizomib is also orally administered and currently in late phase clinical trials. Another strategy could be to use proteasome inhibitors in combination with other agents, e.g. imatinib. Although preliminary results from our laboratory exploring a combination of imatinib and bortezomib did not show clear synergistic effects, alternative administration regimens, such as drug sequencing, could be explored^44^. Combinations with other agents might also be promising. For example, recent strategies include AKT/mTOR inhibitors as bortezomib has been shown to increase AKT activation in multiple myeloma cells^45^.

Further studies will be needed to delineate why compounds that target other regulators of the proteolytic machinery have a significantly lower efficacy in GIST cells than 20S inhibitors. The DUB inhibitor b-AP15 clearly had a lower ability to induce the accumulation of poly-ubiquitinated proteins and resulting H2AX upregulation as well as transcriptional inhibition of *KIT*. This is in line with results from its original report, in which substantially higher concentrations were needed to induce ubiquitination in HCT 116 cells^21^. Any *in vivo* efficacy of this compound will therefore depend on its bioavailability and whether it will be feasible to achieve necessary serum levels paired with acceptable toxicity. Of interest, no clinical trial with b-AP15 has been initiated to this date, and a phase I/II clinical trial testing its derivative, VLX1570^46^, was terminated due to dose-limiting toxicity (clinicaltrials.gov/ct2/show/NCT02372240). Our results for the NAE inhibitor MLN4924 are more complex. The compound exhibited a strong cell cycle inhibitory effect in GIST cells, which correlated well with the demonstrated reduction of ubiquitin-conjugated proteins and its predilection for regulating the turnover of cell cycle-regulating proteins, such as p27^Kip1 27,28^. However, it is not as clear why apoptosis was only induced at much higher concentrations. While MLN4924 has moved forward into several clinical studies, the maximum serum concentrations achieved in these trials were lower than the clinically meaningful dose for GIST, underlining that MLN4924 is not a medication to move forward in this entity^47,48^.

Unfortunately, many preclinical studies never reach clinical success. While the ultimate significance of proteasome inhibitors for the treatment of patients with GIST is difficult to estimate, the fact that our *in vitro* and *in vivo* results are similar to resutls reported in multiple myeloma models is promising. For example, treatment of bortezomib-sensitive multiple myeloma cell lines with delanzomib and bortezomib at concentrations comparable to what we used in our study had a similar effect with respect to the accumulation of ubiquitinated proteins^18^. Another important question for preclinical studies is whether the concentration at which the biologic effects occur are achievable in the serum. Delanzomib serum concentrations for multiple myeloma, for example, while lower than the calculated effective serum concentration needed to treat GIST patients, showed high peak concentrations after initial administration with a rapid distribution into the extra-vascular space^31^. However, the drug was shown to have a prolonged elimination phase and a mean half-life of 62 hours, meaning that drug concentrations in the tumor could be sufficient for exerting a biological effect. This notion is corroborated in our study by the fact that we saw a substantial anti-tumor effect after only one single dose of delanzomib.

Moreover, the effective concentrations determined in our preclinical data for ixazomib are not that different from what has been reported for myeloma models^30^, indicating that this compound might be within therapeutic reach for GIST patients. In contrast, achieving effective serum concentrations of carfilzomib in the patient setting seems difficult.

In summary, we have shown that inhibitors of the ubiquitin-proteasome machinery, especially those that inhibit the 20S proteolytic core, have therapeutic activity in imatinib-sensitive and imatinib-resistant *KIT*-mutant GIST model systems. Clinical studies are encouraged.

## MATERIALS AND METHODS

### Cell culture and inhibitor treatments

An array of imatinib (IM)-sensitive and IM-resistant human GIST cell lines was used in the present study. IM-sensitive cells included GIST882 (kindly provided by Jonathan A. Fletcher, Brigham and Women’s Hospital, Boston, MA, USA)^49^ and GIST-T1 (kindly provided by Takahiro Taguchi, Kochi Medical School, Nankoku Kochi, Japan)^50^. As described previously, GIST882 cells carry a homozygous mutation in *KIT* exon 13 (p.K642E), and GIST-T1 cells are characterized by a heterozygous *KIT* exon 11 deletion (p.V560_Y578del)^50^. The IM-resistant GIST cell lines GIST48 (homozygous *KIT* exon 11 mutation, p.V560D; *KIT* exon 17 point mutation, p.D820A) and GIST430 (heterozygous *KIT* exon 11 deletion, p.V560_L576del; *KIT* exon 13 point mutation, p.V654A) were derived from human GISTs that developed clinical resistance to IM therapy (J.A. Fletcher)^51^. Cells were maintained as described previously^51^.

Cells were treated with carfilzomib (CFZ, Fisher), ixazomib (IXA, SelleckChem), delanzomib (DLZ, SelleckChem, Toronto Research Chemicals), b-AP15 (EMD Millipore) or MLN-4924 (SelleckChem) at the indicated concentrations in DMSO or mock-treated with 0.1% DMSO for up to 72 h, as indicated. Treatment with imatinib mesylate (1 μM in DMSO; LC Laboratories), sunitinib malate (1 μM in DMSO; LC Laboratories), bortezomib (0.01 μM in DMSO; LC Laboratories) as well as α-amanitin (1μg/mL in dH_2_O; Sigma) served as controls.

### Antibodies

Primary antibodies used for immunoblotting, immunofluorescence and immunohistochemistry were actin (Sigma), cleaved caspase 3 (Cell Signaling), CREB binding protein (CBP; Santa Cruz), cyclin A (Novocastra), pH2AX S139 (Millipore), H2AX (Bethyl), phospho-histone H3 S10, phospho-KIT Y719 (both Cell Signaling), Ki-67 (Thermo Scientific Neomarkers), KIT (Dako/Agilent), p27^Kip1^(Fisher/BD Biosciences Pharmingen), PARP (Invitrogen), cleaved PARP (Abcam) and mono-ubiquitin (BD Pharmingen).

### 26S proteasome activity assay

The chymotrypsin-like activity of the proteasome after treatment with the above-mentioned proteasome inhibitors was measured using the luminescence-based Proteasome-Glo assay (Promega). Cells were plated in 96-well flat-bottomed plates (Perkin Elmer), cultured for 24 h and then incubated for 2 h with the respective compounds at indicated concentrations or DMSO-only solvent control. A peptide substrate for the chymotrypsin-like activity of the proteasome conjugated to recombinant luciferase, Suc-LLVY-(succinyl-leucine-leucine-valine-tyrosine-)aminoluciferin, was added for 10 minutes, before luminescence was measured with a BioTek Synergy 2 Luminometer (BioTek). Data were normalized to the DMSO-only control group.

### GIST xenograft models

*In vivo* studies were carried out at the Laboratory of Experimental Oncology, KU Leuven (Leuven, Belgium) according to local guidelines and Belgian regulations and were approved by the Ethical Committee for Laboratory Animal Research, KU Leuven. UZLX-GIST1 (IM-sensitive; *KIT* exon 11 mutation, p.V560D) and UZLX-GIST9 (IM-resistant) were derived from clinical samples of consenting GIST patients treated at the University Hospitals Leuven, as previously described^52,53^. UZLX-GIST9 initially presented with a primary *KIT* exon 11 p.P577del mutation. At time of metastasis, mutational analysis revealed the original mutation as well as a *KIT* exon 11 frameshift (p.W557LfsX5) and a *KIT* exon 13 point mutation (p.D820G)^52^. UZLX-GIST1 (passage 23), UZLX-GIST9 (passage12) and GIST430 (passage 6) xenografts were created by bilateral transplantation of tumor fragments into the flank of female adult athymic nude mice (NMRI, nu/nu; Janvier Laboratories). When tumors were 1 cm in diameter, mice were randomized into groups of 4-6 animals each for different treatment regimens. Delanzomib (in 5% mannitol:propylene glycol [4:1 v/v]) was administered by oral gavage twice a week for three weeks. IM-treated mice (50 mg/kg twice a day, p.o.) served as positive control, while mice receiving vehicle were used as negative control. A maximum-tolerated dose experiment for delanzomib using 6 *vs.* 8 mg/kg was conducted. All mice tolerated the higher dose of 8 mg/kg well (data not shown), which was used in all experiments. Above doses were chosen according to the literature, where oral dosing between 7.8 mg/kg and 13 mg/kg was tested in myeloma models^18,54^. Our selected dose of 8 mg/kg was at the low end of the spectrum for myeloma studies and should hence well translate into the human setting. Tumor volume, weight and general health of the mice were recorded^52^. After the mice were sacrificed, tumors were excised and divided for fresh frozen samples and histopathologic examination. Histopathologic grading of the response to the compounds was based on the microscopic amount of necrosis, myxoid degeneration or fibrosis with grade 1 representing a minimal response (0-10%) and grade 4 representing a maximal response (>90%)^55^.

### Statistics

Statistical significance was assessed using Student’s t test for independent samples. *P* values ≤0.05 were considered significant.

## Supplemental Methods

Further methods used are listed under Supplemental Information.

## DATA AVAILABILITY

The data set generated during and/or analyzed during the current study are available from the corresponding author on reasonable request.

## ACKNOWLEDGEMENTS

The authors would like to thank Jonathan A. Fletcher and Takahiro Taguchi for sharing important reagents, Sarangarajan Ranganathan and Ivy John for expertise in mouse histopathology, Jennifer D. Francis, Laura Presutti, Eric W. Weil and Michelle Wolf for technical assistance as well as Stefan Duensing for critically reading the manuscript and helpful discussions. This work was supported by a Research Scholar Grant from the American Cancer Society (RSG-08-092-01-CCG; to A.D), the GIST Cancer Research Fund (to A.D.), The Life Raft Group (to A.D. and M.D.R.), the Out of the Woods Foundation (to A.D.), the Flemish Cancer Society (to P.S.) and numerous private donations (to A.D.). A.D. is supported by the UPMC Hillman Cancer Center and in part by a grant from the Pennsylvania Department of Health. The Department specifically disclaims responsibility for any analyses, interpretations or conclusions.

## AUTHOR CONTRIBUTIONS

Conception and design: AD. Development of methodology: JLR, AAA, KRMe, KRMa, MDR, AW, AD. Acquisition of data: JLR, AAA, DML, YKG, KRMe, AA, SSP, YT, JW, HSD, MDR, AW, AD. Analysis and interpretation of data: JLR, AAA, DML, KRMe, AA, SSP, AW, AD. Writing, review and/or revision of the manuscript: JLR, DML, SSP, MDR, AW, AD. Administrative, technical, or material support: KRMa, PS, MDR, AD. Study supervision: AD.

## REFERENCES

1. Verweij, J. et al. Progression-free survival in gastrointestinal stromal tumours with high-dose imatinib: randomised trial. Lancet 364, 1127–1134 (2004).

2. Gramza, A. W., Corless, C. L. & Heinrich, M. C. Resistance to Tyrosine Kinase Inhibitors in Gastrointestinal Stromal Tumors. Clin Cancer Res 15, 7510–7518 (2009).

3. Voges, D., Zwickl, P. & Baumeister, W. The 26S proteasome: A molecular machine designed for controlled proteolysis. Annu Rev Biochem 68, 1015–1068 (1999).

4. Manasanch, E. E. & Orlowski, R. Z. Proteasome inhibitors in cancer therapy. Nat. Publ. Gr. 1–17 (2017). doi:10.1038/nrclinonc.2016.206

5. Bauer, S. et al. Proapoptotic activity of bortezomib in gastrointestinal stromal tumor cells. Cancer Res. 70, 150–159 (2010).

6. Liu, Y. et al. Histone H2AX is a mediator of gastrointestinal stromal tumor cell apoptosis following treatment with imatinib mesylate. Cancer Res. 67, 2685–2692 (2007).

7. Liu, Y., Parry, J. A. J. A., Chin, A., Duensing, S. & Duensing, A. Soluble histone H2AX is induced by DNA replication stress and sensitizes cells to undergo apoptosis. Mol. Cancer 7, 61 (2008).

8. Rausch, J. L. et al. Opposing roles of KIT and ABL1 in the therapeutic response of gastrointestinal stromal tumor (GIST) cells to imatinib mesylate. Oncotarget 8, 4471–4483 (2017).

9. Maki, R. G. et al. A multicenter Phase II study of bortezomib in recurrent or metastatic sarcomas. Cancer 103, 1431–1438 (2005).

10. Bahleda, R. et al. Phase I trial of bortezomib daily dose: safety, pharmacokinetic profile, biological effects and early clinical evaluation in patients with advanced solid tumors. 1–10 (2017). doi:10.1007/s10637-017-0531-3

11. Deming, D. A. et al. A Phase I study of intermittently dosed vorinostat in combination with bortezomib in patients with advanced solid tumors. Invest. New Drugs 32, 323–329 (2013).

12. Papandreou, C. N. et al. Phase I Trial of the Proteasome Inhibitor Bortezomib in Patients With Advanced Solid Tumors With Observations in Androgen-Independent Prostate Cancer. J. Clin. Oncol. 22, 2108–2121 (2004).

13. Demo, S. D. et al. Antitumor activity of PR-171, a novel irreversible inhibitor of the proteasome. Cancer Res. 67, 6383–6391 (2007).

14. Herndon, T. M. et al. U.s. Food and Drug Administration approval: carfilzomib for the treatment of multiple myeloma. Clin Cancer Res 19, 4559–4563 (2013).

15. O’Connor, O. A. et al. A phase 1 dose escalation study of the safety and pharmacokinetics of the novel proteasome inhibitor carfilzomib (PR-171) in patients with hematologic malignancies. Clin Cancer Res 15, 7085–7091 (2009).

16. Kupperman, E. et al. Evaluation of the proteasome inhibitor MLN9708 in preclinical models of human cancer. Cancer Res. 70, 1970–1980 (2010).

17. Shirley, M. Ixazomib: First Global Approval. Drugs 76, 405–411 (2016).

18. Piva, R. et al. CEP-18770: A novel, orally active proteasome inhibitor with a tumor-selective pharmacologic profile competitive with bortezomib. Blood 111, 2765–2775 (2008).

19. Dorsey, B. D. et al. Discovery of a potent, selective, and orally active proteasome inhibitor for the treatment of cancer. J. Med. Chem. 51, 1068–1072 (2008).

20. Vogl, D. T. et al. Phase I/II study of the novel proteasome inhibitor delanzomib (CEP-18770) for relapsed and refractory multiple myeloma. Leuk. & Lymphoma 58, 1872–1879 (2016).

21. D’Arcy, P. aacute draig et al. Inhibition of proteasome deubiquitinating activity as a new cancer therapy. Nat. Med. 17, 1636–1640 (2011).

22. Oh, Y.-T., Deng, L., Deng, J. & Sun, S.-Y. The proteasome deubiquitinase inhibitor b-AP15 enhances DR5 activation-induced apoptosis through stabilizing DR5. Sci. Rep. 1–11 (2017). doi:10.1038/s41598-017-08424-w

23. Tian, Z. et al. A novel small molecule inhibitor of deubiquitylating enzyme USP14 and UCHL5 induces apoptosis in multiple myeloma and overcomes bortezomib resistance. Blood 123, 706–716 (2014).

24. Chitta, K. et al. Targeted inhibition of the deubiquitinating enzymes, USP14 and UCHL5, induces proteotoxic stress and apoptosis in Waldenström macroglobulinaemia tumour cells. Br. J. Haematol. 169, 377–390 (2015).

25. Kumar, S., Yoshida, Y. & Noda, M. Cloning of a cDNA Which Encodes a Novel Ubiquitin-like Protein. Biochem. Biophys. Res. Commun. 195, 393–399 (1993).

26. Hori, T. et al. Covalent modification of all members of human cullin family proteins by NEDD8. Oncogene 18, 6829–6834 (1999).

27. Soucy, T. A., Dick, L. R., Smith, P. G., Milhollen, M. A. & Brownell, J. E. The NEDD8 conjugation pathway and its relevance in cancer biology and therapy. Genes & cancer 1, 708–716 (2010).

28. Soucy, T. A. et al. An inhibitor of NEDD8-activating enzyme as a new approach to treat cancer. Nature 458, 732–736 (2009).

29. Chauhan, D. et al. A small molecule inhibitor of ubiquitin-specific protease-7 induces apoptosis in multiple myeloma cells and overcomes bortezomib resistance. Cancer Cell 22, 345–358 (2012).

30. Chauhan, D. et al. In vitro and in vivo selective antitumor activity of a novel orally bioavailable proteasome inhibitor MLN9708 against multiple myeloma cells. Clin Cancer Res 17, 5311–5321 (2011).

31. Gallerani, E. et al. A first in human phase I study of the proteasome inhibitor CEP-18770 in patients with advanced solid tumours and multiple myeloma. Eur J Cancer 49, 290–296 (2013).

32. Li, H. et al. Pazopanib, a Receptor Tyrosine Kinase Inhibitor, Suppresses Tumor Growth through Angiogenesis in Dedifferentiated Liposarcoma Xenograft Models. Transl. Oncol. 7, 665–671 (2014).

33. Floris, G. et al. High Efficacy of Panobinostat Towards Human Gastrointestinal Stromal Tumors in a Xenograft Mouse Model. Clin. Cancer Res. 15, 4066–4076 2009).

34. Liu, Y. et al. Imatinib mesylate induces quiescence in gastrointestinal stromal tumor cells through the CDH1-SKP2-p27Kip1 signaling axis. Cancer Res. 68, 9015–9023 (2008).

35. Boichuk, S. et al. The DREAM complex mediates GIST cell quiescence and is a novel therapeutic target to enhance imatinib-induced apoptosis. Cancer Res. 73, 5120–5129 (2013).

36. Geng, F., Wenzel, S. & Tansey, W. P. Ubiquitin and Proteasomes in Transcription. Annu Rev Biochem 81, 177–201 (2012).

37. Durairaj, G. & Kaiser, P. The 26S Proteasome and Initiation of Gene Transcription. Biomolecules 4, 827–847 (2014).

38. Chi, P. et al. ETV1 is a lineage survival factor that cooperates with KIT in gastrointestinal stromal tumours. Nature 467, 849–853 (2010).

39. Park, G. H., Plummer, H. K. & Krystal, G. W. Selective Sp1 binding is critical for maximal activity of the human c-kit promoter. Blood 92, 4138–4149 (1998).

40. Karki, K., Harishchandra, S. & Safe, S. Bortezomib Targets Sp Transcription Factors in Cancer Cells. Mol. Pharmacol. 94, 1187–1196 (2018).

41. Koyama, D. et al. Proteasome inhibitors exert cytotoxicity and increase chemosensitivity via transcriptional repression of Notch1 in T-cell acute lymphoblastic leukemia. Leukemia 28, 1216–1226 (2014).

42. Britton, M. et al. Selective Inhibitor of Proteasome• s Caspase-like Sites Sensitizes Cells to Specific Inhibition of Chymotrypsin-like Sites. Chem. Biol. 16, 1278–1289 (2009).

43. Chauhan, D. et al. A novel orally active proteasome inhibitor induces apoptosis in multiple myeloma cells with mechanisms distinct from Bortezomib. Cancer Cell 8, 407–419 (2005).

44. Serrano, C. et al. Complementary activity of tyrosine kinase inhibitors against secondary kit mutations in imatinib-resistant gastrointestinal stromal tumours. Br. J. Cancer 120, (2019).

45. Kapoor, P., Ramakrishnan, V. & Rajkumar, S. V. Bortezomib Combination Therapy in Multiple Myeloma. Semin. Hematol. 49, 228–242 (2012).

46. Wang, X. et al. The proteasome deubiquitinase inhibitor VLX1570 shows selectivity for ubiquitin-specific protease-14 and induces apoptosis of multiple myeloma cells. Sci. Rep. 6, 26979 (2016).

47. Shah, J. J. et al. Phase I Study of the Novel Investigational NEDD8-Activating Enzyme Inhibitor Pevonedistat (MLN4924) in Patients with Relapsed/Refractory Multiple Myeloma or Lymphoma. Clin Cancer Res 22, 34–43 (2016).

48. Bhatia, S. et al. A phase I study of the investigational NEDD8-activating enzyme inhibitor pevonedistat (TAK-924/MLN4924) in patients with metastatic melanoma. Invest. New Drugs 34, 439–449 (2016).

49. Duensing, A. et al. Mechanisms of oncogenic KIT signal transduction in primary gastrointestinal stromal tumors (GISTs). Oncogene 23, 3999–4006 (2004).

50. Taguchi, T. et al. Conventional and molecular cytogenetic characterization of a new human cell line, GIST-T1, established from gastrointestinal stromal tumor. Lab. Invest. 82, 663–665 (2002).

51. Boichuk, S. et al. Unbiased compound screening identifies unexpected drug sensitivities and novel treatment options for gastrointestinal stromal tumors. Cancer Res. 74, 1200–1213 (2014).

52. Van Looy, T. et al. Characterization and assessment of the sensitivity and resistance of a newly established human gastrointestinal stromal tumour xenograft model to treatment with tyrosine kinase inhibitors. Clin. Sarcoma Res. 4, 10 (2014).

53. Van Looy, T. et al. Therapeutic Efficacy Assessment of CK6, a Monoclonal KIT Antibody, in a Panel of Gastrointestinal Stromal Tumor Xenograft Models. Transl. Oncol. 8, 112–118 (2015).

54. Sanchez, E. et al. The proteasome inhibitor CEP-18770 enhances the anti-myeloma activity of bortezomib and melphalan. Br. J. Haematol. 148, 569–581 (2010).

55. Agaram, N. P. et al. Pathologic and molecular heterogeneity in imatinib-stable or imatinib-responsive gastrointestinal stromal tumors. Clin Cancer Res 13, 170–181 (2007).

