## Supplemental text and figures for "Differential antitumor activity of compounds targeting the ubiquitin-proteasome machinery in gastrointestinal stromal tumor (GIST) cells"

### SUPPLEMENTARY METHODS

#### *Immunological and cell staining methods*

Protein lysates of cells growing as monolayer were prepared by scraping cells into lysis buffer (1% NP-40, 50 mM Tris-HCl pH 8.0, 100 mM sodium fluoride, 30 mM sodium pyrophosphate, 2 mM sodium molybdate, 5 mM EDTA, 2 mM sodium orthovanadate) containing protease inhibitors (10 µg/ml aprotinin, 10 µg/ml leupeptin, 1 µM phenylmethylsulfonyl fluoride). Lysates were incubated for 1 h with shaking at 4°C and then cleared by centrifugation for 30 min at 14,000 rpm (4°C). Protein concentrations were determined using the Bradford assay (Biorad). Thirty µg of protein were loaded on a 4-12% Bis-Tris gel (Invitrogen) and blotted onto a nitrocellulose membrane.

For immunofluorescence analysis, cells grown in chamber slides (BD Biosciences) were briefly washed in PBS and fixed in 4% paraformaldehyde in PBS for 15 min at room temperature (RT). Cells were then washed in PBS and permeabilized with 1% Triton-X 100 in PBS for 15 min (RT) followed by washing in PBS and blocking with 10% normal donkey serum (Jackson Immunoresearch) for 15 min. Primary antibody incubation was done overnight at 4°C in a humidified chamber and an additional hour at 37°C the next morning. After a brief wash in PBS, cells were incubated with Alexa Fluor 488-conjugated goat anti-rabbit secondary antibodies (Invitrogen; 30 min, RT), washed (PBS) and counterstained with 4',6-diamidino-2-phenylindole (DAPI; Vector Laboratories). Cells were analyzed using an Olympus AX70 epifluorescence microscope equipped with a SPOT RT digital camera and SPOT imaging software.

Immunohistochemistry was performed on sections (4µm) of formalin-fixed, paraffin-embedded mouse tumors from *in vivo* experiments. Slides were deparaffinized, underwent antigen retrieval and were incubated with rabbit monoclonal primary antibodies. The signal was detected using anti-rabbit EnVision+ System-HRP secondary antibodies and 3'diaminobenzidine-tetrahydrochloride (DAB; both from Dako/Agilent). Apoptotic and mitotic cells were visualized by immunohistochemical staining for cleaved PARP and phosphorylated histone H3 S10, respectively, and Ki-67 was used to evaluate cellular proliferation[35]. The number of positive cells was counted per 10 high power fields (HFP) at 400-fold magnification.

Detection of apoptotic cells *in vitro* was done using the In situ Cell Death Detection Kit (Roche Applied Sciences) according to manufacturer's recommendations[5].

***Luminescence-based proliferation and apoptosis assays***

Cell viability and apoptosis studies were performed using the CellTiter-Glo (readout: ATP) and Caspase-Glo (readout: caspase 3/7 activity) luminescence-based assays (Promega). Cells were plated in 96-well flat-bottomed plates (Perkin Elmer), cultured for 24 h and then incubated for 48 h (Caspase-Glo) or 72 h (CellTiter-Glo) with the respective compounds at indicated concentrations or DMSO-only solvent control. Luminescence was measured with a BioTek Synergy 2 Luminometer (BioTek). Data were normalized to the DMSO-only control group.

***Reverse transcriptase (RT)-PCR and quantitative real time RT-PCR***

RT-PCR and quantitative real time RT-PCR (qRT-PCR) were performed as described previously[5]. Exon-overlapping, mRNA/cDNA-specific primers were used to amplify *KIT* (forward: 5'-TCATGGTCGGATCACAAAGA-3', reverse: 5'-AGGGGCTGCTTCCTAAAGAG-3'; Operon) and *β-actin* (forward: 5'-CCAAGGCCAACCGCGAGAAGATGAC-3', reverse: 5'-AGGGTACATGGTGGTGCCGCCAGAC-3'). *β-actin* served as reference gene for relative quantification in quantitative real time RT-PCR (qRT-PCR) experiments.

### SUPPLEMENTARY FIGURE LEGENDS

**Supplementary Figure S1. Delanzomib is highly effective in IM-sensitive and IM-resistant GIST cells and rapidly leads to a dose- and time-dependent accumulation of ubiquitinated proteins.**

**(A)** Immunoblot analysis for markers of cell cycle regulation and apoptosis after treatment with increasing concentrations (0.0001 μM – 10 μM) of delanzomib (DLZ). Grouped immunoblot images are either cropped from different parts of the same gel or from a separate gel run with another aliquot of the same protein lysate.

**(B)** Dose-dependent effect of delanzomib on induction of apoptosis in IM-sensitive GIST-T1 and IM-resistant GIST430 cells as measured by TUNEL assay. Graphs represent mean and standard error of at least three experiments with at least 100 cells counted each. **, $p \leq 0.01$ in comparison to control; ***, $p \leq 0.001$ in comparison to control (Student's t-test, 2-tailed).

**(C)** Immunoblot analysis for markers of cell cycle regulation and apoptosis after treatment of IM-sensitive (GIST882) and IM-resistant (GIST430) cells with DMSO or delanzomib (0.1 μM) for the indicated times.

**(D, E)** Dose- **(D)** and time-dependent **(E)** accumulation of mono-ubiquitinated proteins in IM-sensitive (GIST-T1) and IM-resistant (GIST48) cells after delanzomib treatment as determined by immunoblotting. As expected, kinase inhibitor treatment with imatinib (IM) or sunitinib (SU; both 1.0 μM) did not lead to increased ubiquitination when compared to DMSO-treated controls. IM and SU serve as standard treatment controls for IM-naïve and IM-resistant cell lines, respectively. Bortezomib (BO) serves as control for proteasome inhibition.

**Supplementary Figure S2. Carfilzomib (CFZ) and ixazomib (IXA) lead to cell cycle exit and apoptosis in IM-sensitive and IM-resistant GIST cells.**

**(A–C)** Dose-dependent effect of carfilzomib and ixazomib on cell cycle exit and induction of apoptosis as measured by immunoblotting **(A)** and TUNEL assay **(B, C)**. Graphs represent mean and standard error of at least three experiments with at least 100 cells counted each. *, $p \leq 0.05$ in comparison to control; **, $p \leq 0.01$ in comparison to control; ***, $p \leq 0.001$ in comparison to control (Student's t-test, 2-tailed).

**(D, E)** Time-dependent effect of carfilzomib **(D)** and ixazomib **(E)** on cell cycle exit and induction of apoptosis of GIST cells as determined by immunoblotting. Treatment with carfilzomib (1.0 µM) **(D)** or ixazomib (0.1 µM) **(E)**, respectively, for the indicated times.

**(A, D, E)** Grouped immunoblot images are either cropped from different parts of the same gel or from a separate gel run with another aliquot of the same protein lysate.

**Supplementary Figure S3. Carfilzomib (CFZ) and ixazomib (IXA) lead to a dose- and time-dependent accumulation of ubiquitinated proteins, but to a lesser extent than delanzomib.**

**(A, B)** Dose- **(A)** and time-dependent **(B)** accumulation of mono-ubiquitinated proteins in GIST cells after carfilzomib treatment (0.0001 µM – 10 µM, **(A)**; 1.0 µM **(B)**) as determined by immunoblotting. As expected, kinase inhibitor treatment with IM or SU (both 1.0 µM) did not lead to increased ubiquitination when compared to DMSO-treated controls. Note that the effect seems to be transient and early effects may have been missed in the 72 h-treated dose-response shown in **(A)**. IM and SU serve as standard treatment controls for IM-naïve and IM-resistant cell lines, respectively. Bortezomib (BO) serves as control for proteasome inhibition.

**(C, D)** Dose- **(C)** and time-dependent **(D)** accumulation of mono-ubiquitinated proteins in GIST cells after ixazomib treatment (0.0001 μM – 10 μM, **(C)**; 0.1 μM **(D)**) as determined by immunoblotting. As expected, kinase inhibitor treatment with IM or SU (both 1.0 μM; 72 h) did not lead to increased ubiquitination when compared to DMSO-treated controls.

**Supplementary Figure S4. The DUB inhibitor b-AP15 and the NAE inhibitor MLN4924 are not as effective as 26S proteasome inhibitors.**

**(A, B)** Immunoblot analysis for markers of cell cycle regulation and apoptosis in GIST cells after treatment with increasing concentrations (0.0001 μM – 10 μM; 72 h) of b-AP15 **(A)** or MLN4924 **(B)**. Treatment with IM or SU (both 1.0 μM) serve as standard treatment controls for IM-naïve and IM-resistant cell lines, respectively. Grouped immunoblot images are either cropped from different parts of the same gel or from a separate gel run with another aliquot of the same protein lysate.

**Supplementary Figure S5. Delanzomib (DLZ) treatment leads to upregulation of soluble histone H2AX and transcriptional downregulation of KIT expression.**

**(A, B)** Immunoblot analysis of GIST-T1 and GIST430 cells treated with delanzomib at the indicated concentrations for 72 h **(A)** or with 0.1 μM delanzomib for the indicated times **(B)** and probed for phospho-H2AX (S139) and total H2AX as well as phospho-KIT (Y719) and total KIT. Grouped immunoblot images are either cropped from different parts of the same gel or from a separate gel run with another aliquot of the same protein lysate.

**(C)** Immunofluorescence microscopic analysis of GIST-T1 and GIST430 cells treated with DMSO or 0.1 μM delanzomib for 72 h and stained for the transcriptional co-activator CREB-binding protein (CBP; green). Nuclei were stained with DAPI. Scale bar, 20 μm.

**Supplementary Figure S6. Carfilzomib and ixazomib lead to upregulation of soluble histone H2AX and transcriptional downregulation of KIT expression, similar to delanzomib.**

**(A – D)** Immunoblot analysis of GIST cells treated with carfilzomib **(A, B)** or ixazomib **(C, D)** at the indicated concentrations for 72 h **(A, C)** or with 1.0 µM carfilzomib **(B)** or with 0.1 µM ixazomib **(D)** for the indicated times and probed for phospho-H2AX (S139) and total H2AX as well as phospho-KIT (Y719) and total KIT.

**(E, F)** Immunofluorescence microscopic analysis of GIST cells treated with DMSO or 1.0 µM carfilzomib **(E)** or 0.1 µM ixazomib **(F)** for 48 h and stained for the transcriptional co-activator CREB-binding protein (CBP; green). Nuclei were stained with DAPI. Scale bar, 20 µm.

**(A – D)** Grouped immunoblot images are either cropped from different parts of the same gel or from a separate gel run with another aliquot of the same protein lysate.

**Supplementary Figure S7. Re-distribution of transcriptional co-factor CREB binding protein (CBP) staining after delanzomib treatment over time.**

Immunofluorescence microscopic analysis of GIST882 cells treated with DMSO or 0.1 µM delanzomib for up to 72 h and stained for CBP (green). Nuclei were stained with DAPI. Note CBP redistribution into large nuclear displacement foci starting at 24 h and complete loss at later time points. Scale bar, 20 µm.

**Supplementary Figure S8. Delanzomib has *in vivo* antitumor activity in GIST xenografts.**

**(A)** Histopathologic and immunohistochemical response of imatinib-resistant UZLX-GIST9 xenografts after 24 h to a one-time bolus treatment with delanzomib in comparison with placebo. Hematoxylin and eosin (H&E) as well as immunohistochemical staining for Ki-67 or cleaved PARP (10X magnification). Histopathologic response grade was determined according to Agaram et al.[55]. Data shown in graph represent at least ten tumors per group. Quantification of proliferation index (% Ki-67 positive cells) and percentage of cleaved PARP-positive cells (brown staining, respectively) represents the average of at least nine tumors per group. Columns, mean + SE; *, p≤0.05, **, p≤0.01 in comparison to control (Student's t-test, 2-tailed).

**(B)** Waterfall plot of best response of percent change of UZLX-GIST1 xenografts during a 21-day treatment with vehicle (black bars), delanzomib (white bars) or imatinib (grey bars).

**(C)** Spider plot of relative tumor volume of GIST430 xenografts treatment with vehicle or delanzomib over the course of a 21-day. *Placebo*, black lines; *delanzomib*, grey dotted lines.

**C)** Histopathologic response of imatinib-resistant GIST430 xenografts to treatment with delanzomib in comparison with placebo. Histopathologic response grade was determined according to Agaram et al.[55]. Data shown in graph represent at least eight tumors per group. Columns, mean + SE; *, p≤0.05 in comparison to control (Student's t-test, 1-tailed).

**(D)** Reduced mitotic activity of GIST430 xenografts after treatment with delanzomib in comparison with placebo. Quantification of mitotic cells (pH3-positive cells per 10 HPF; brown staining) represents the average of at least eight tumors per group. Columns, mean + SE; ***, p≤0.001 in comparison to control (Student's t-test, 2-tailed).

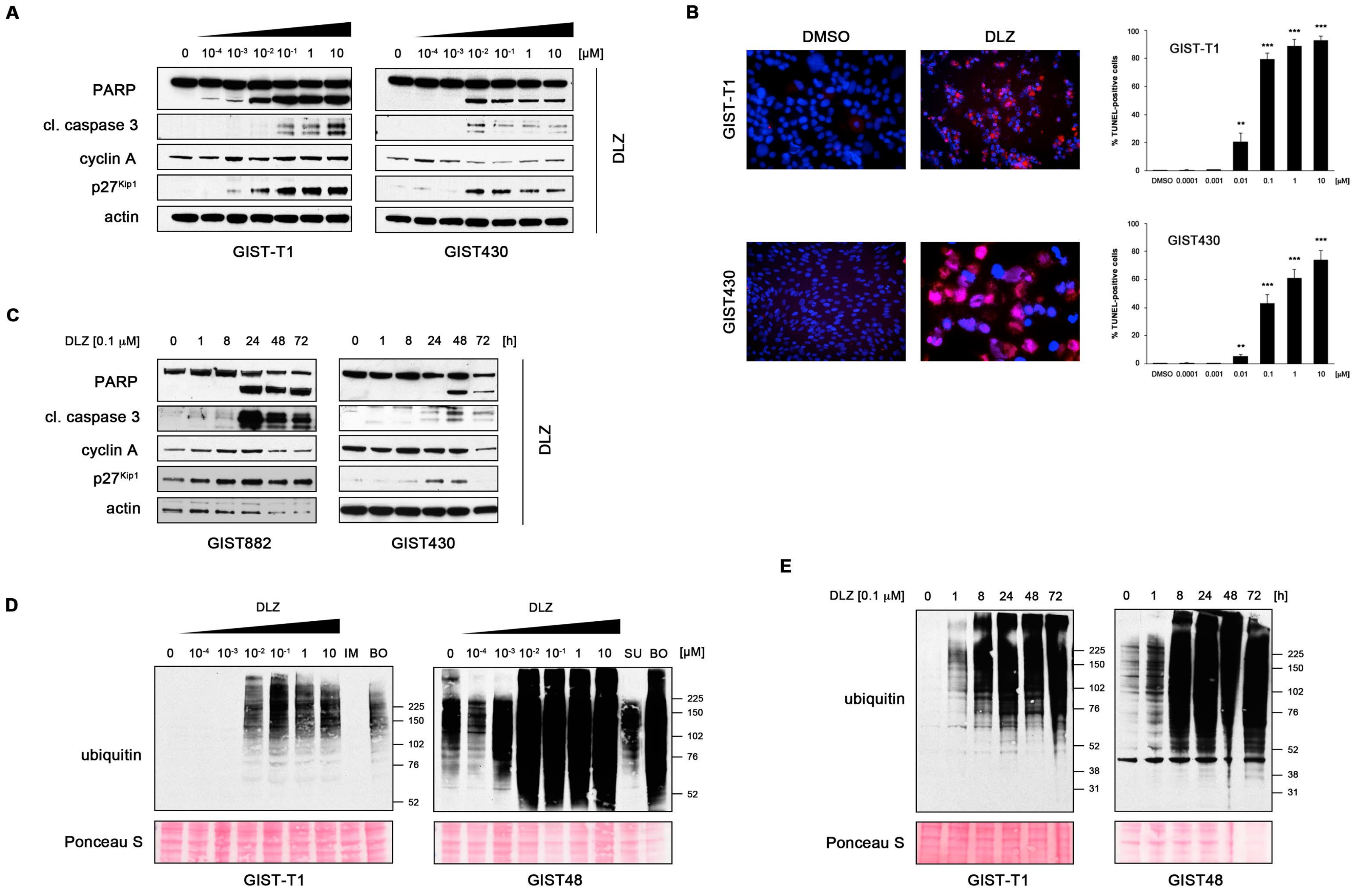


Rausch *et al.*, Supplemental Figure 1

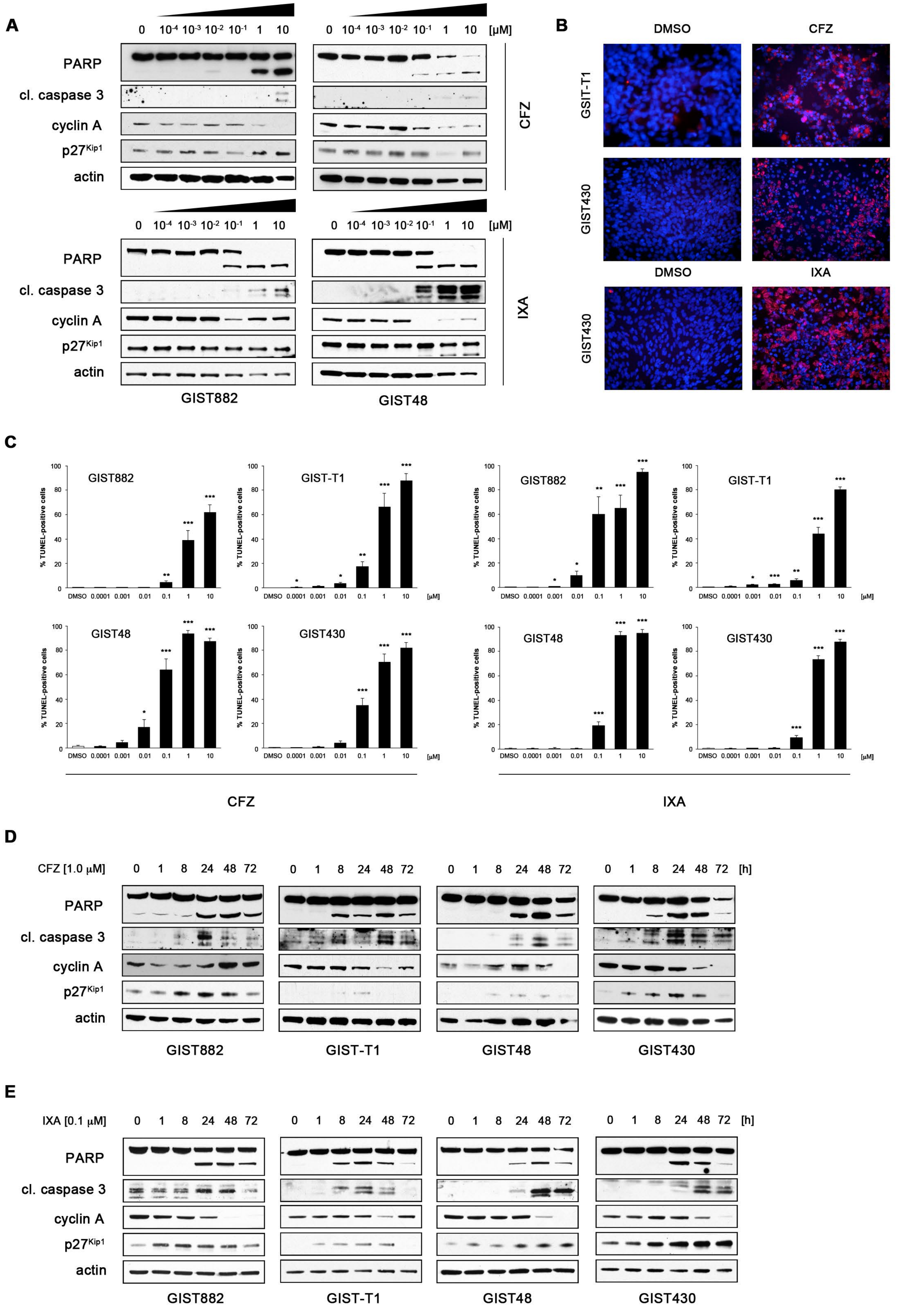


Rausch *et al.*, Supplemental Figure 2

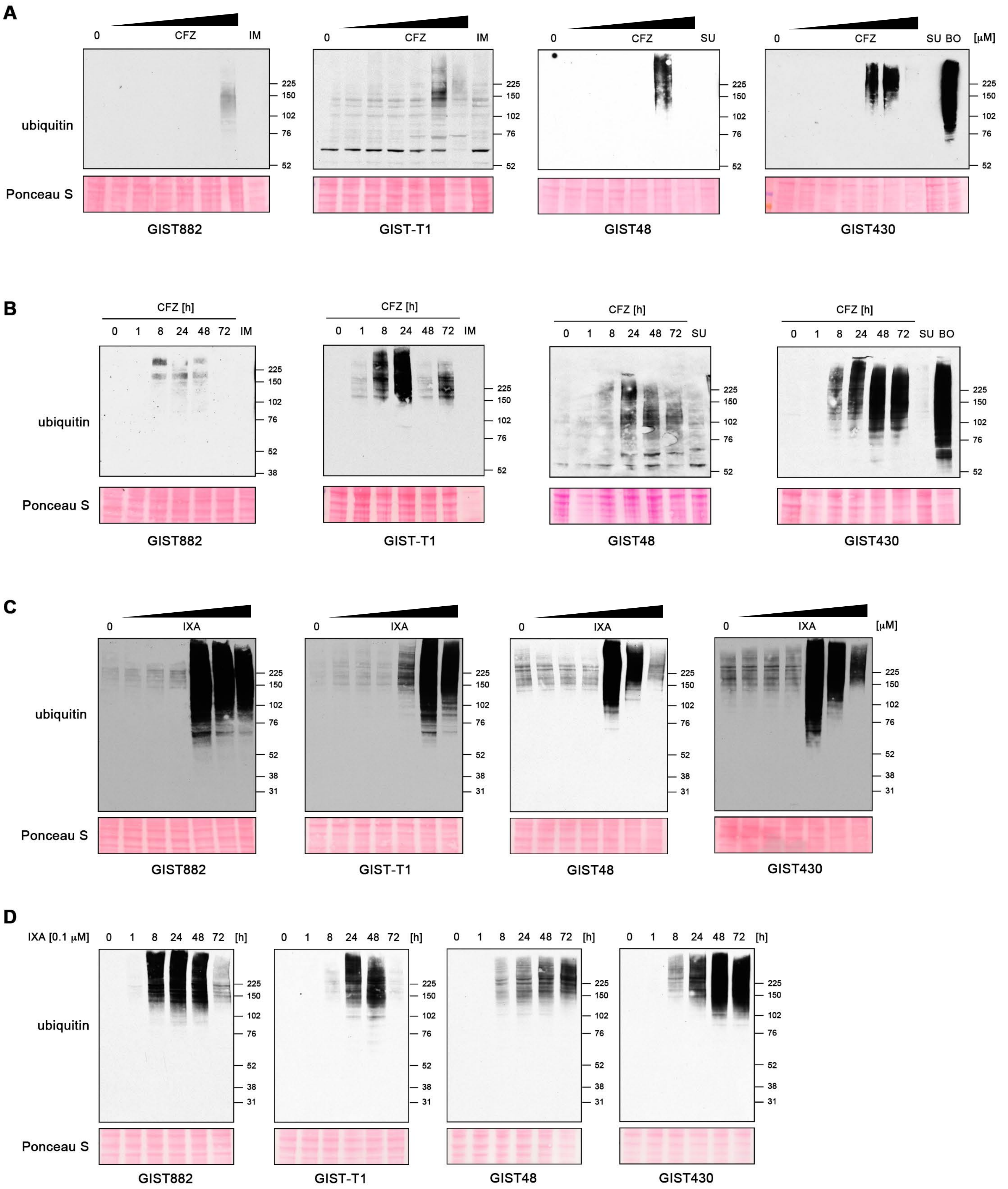


Rausch *et al.*, Supplemental Figure 3

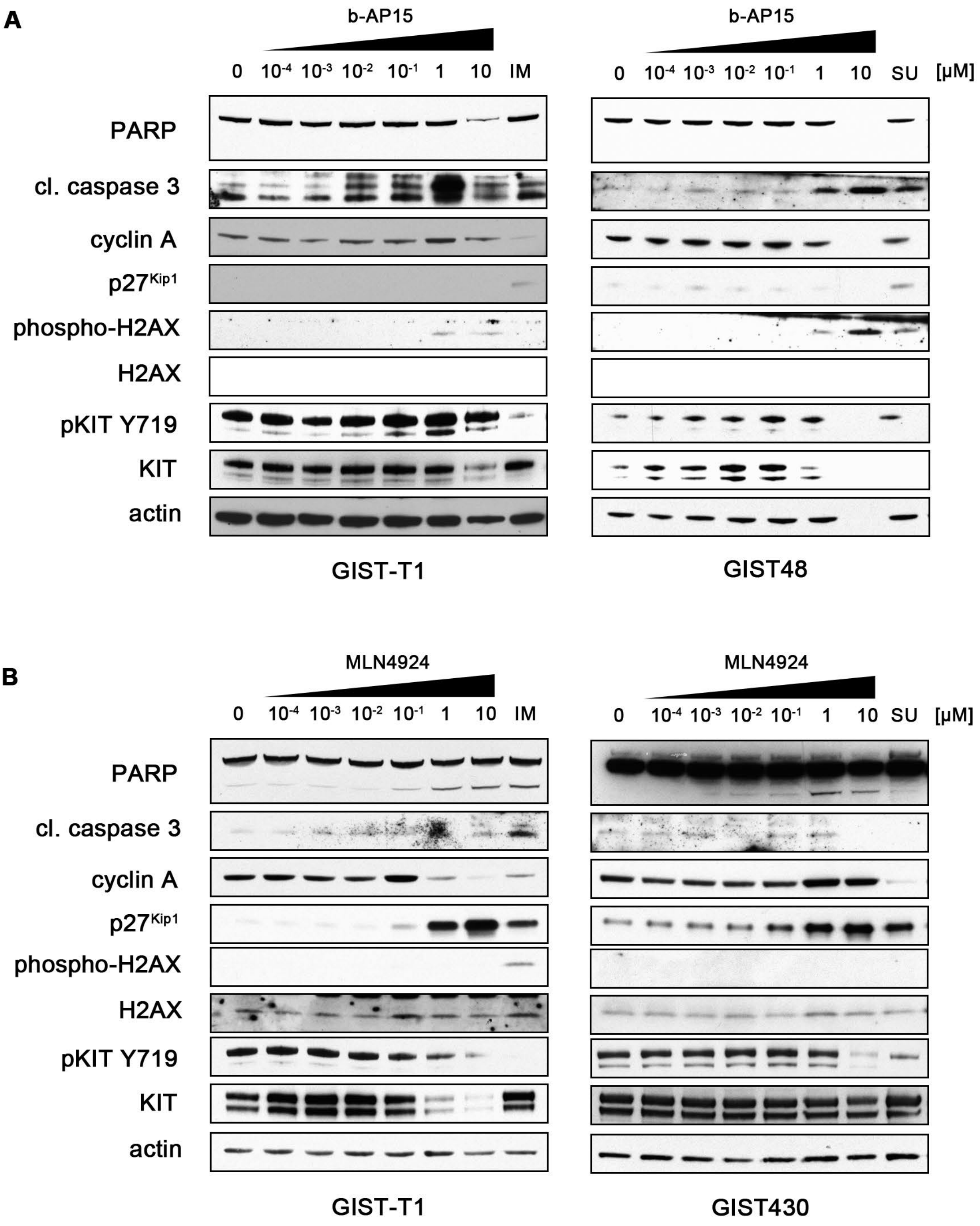


Rausch *et al.*, Supplemental Figure 4

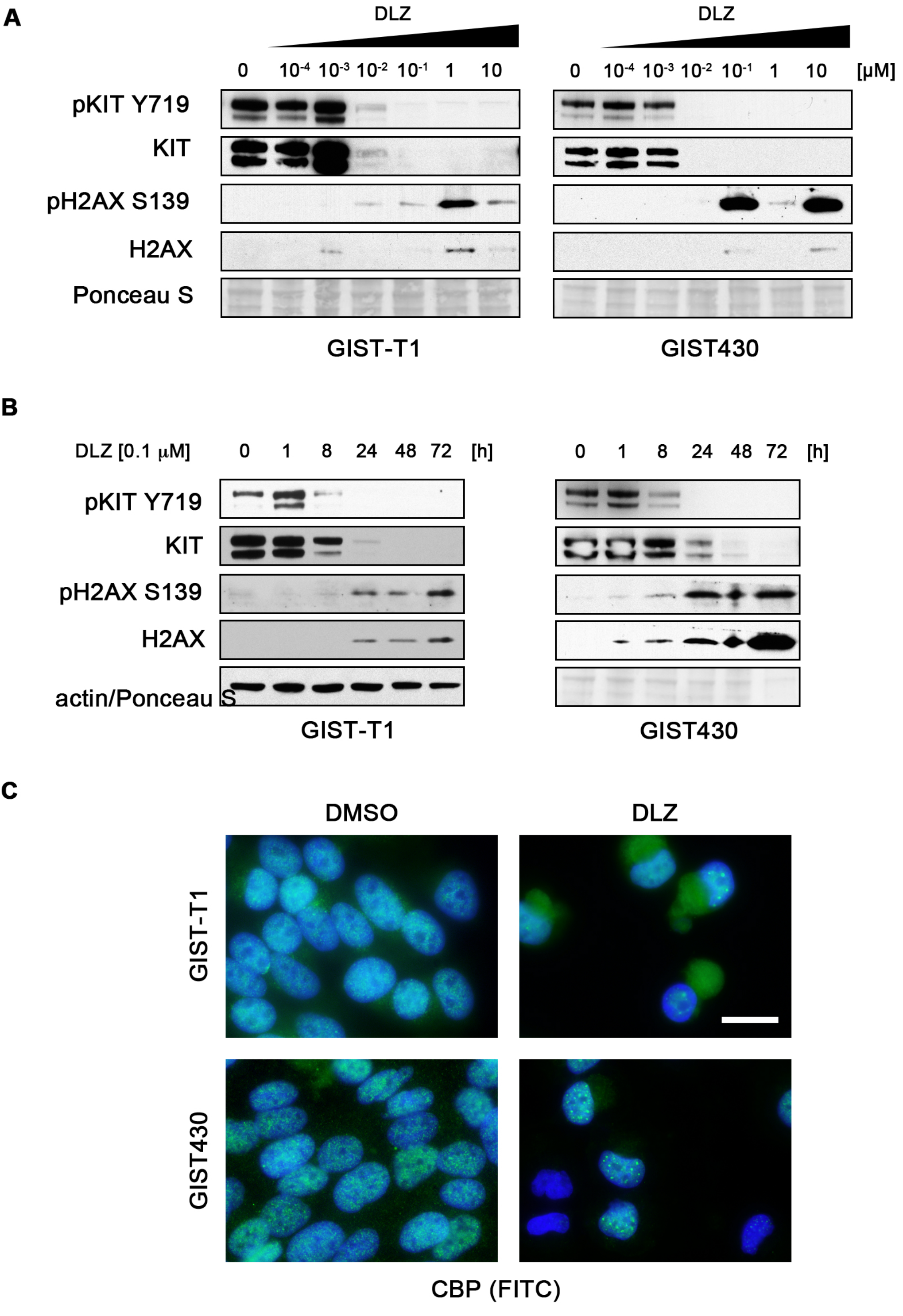


Rausch *et al.*, Supplemental Figure 5

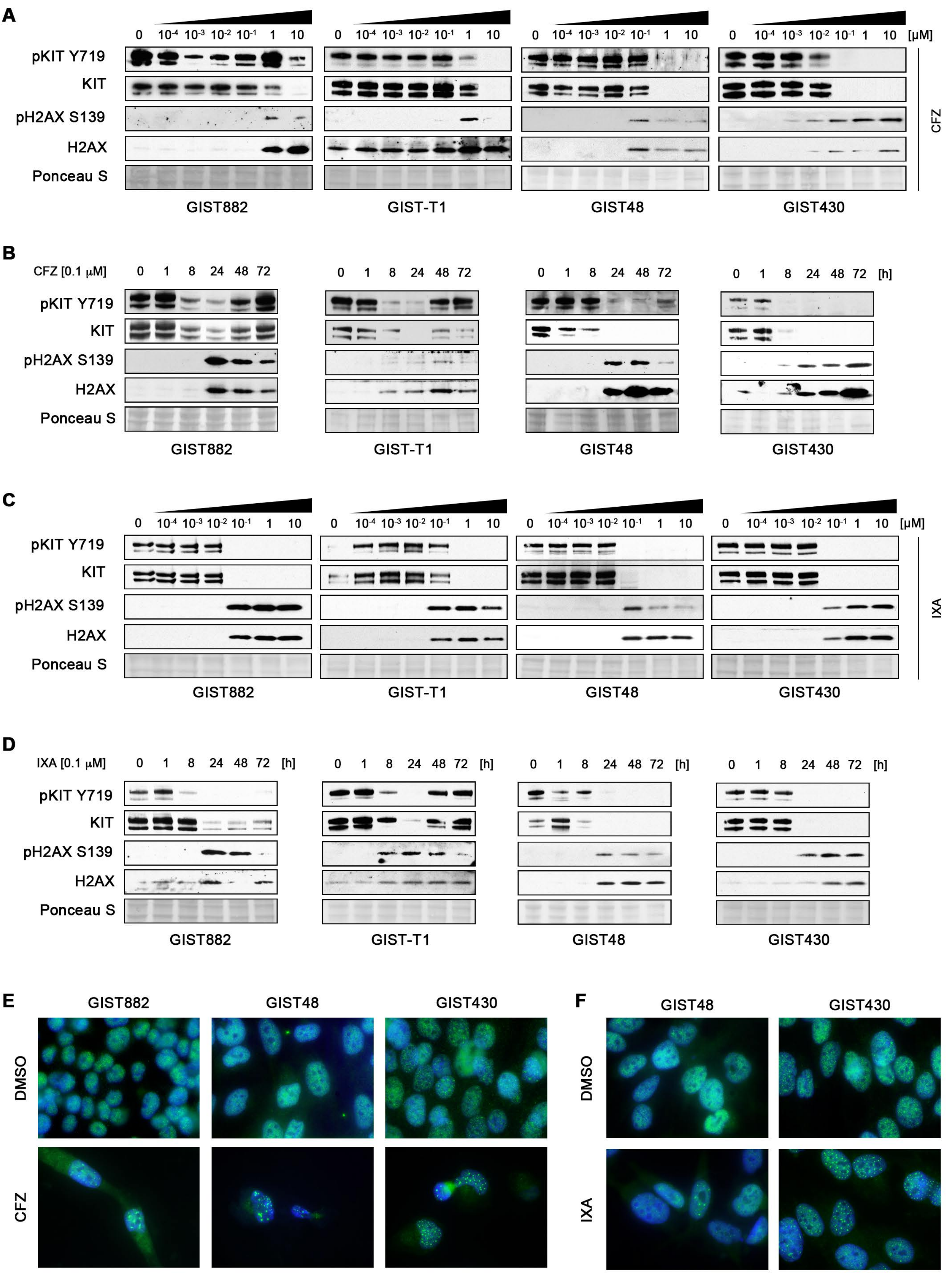


Rausch *et al.*, Supplemental Figure 6

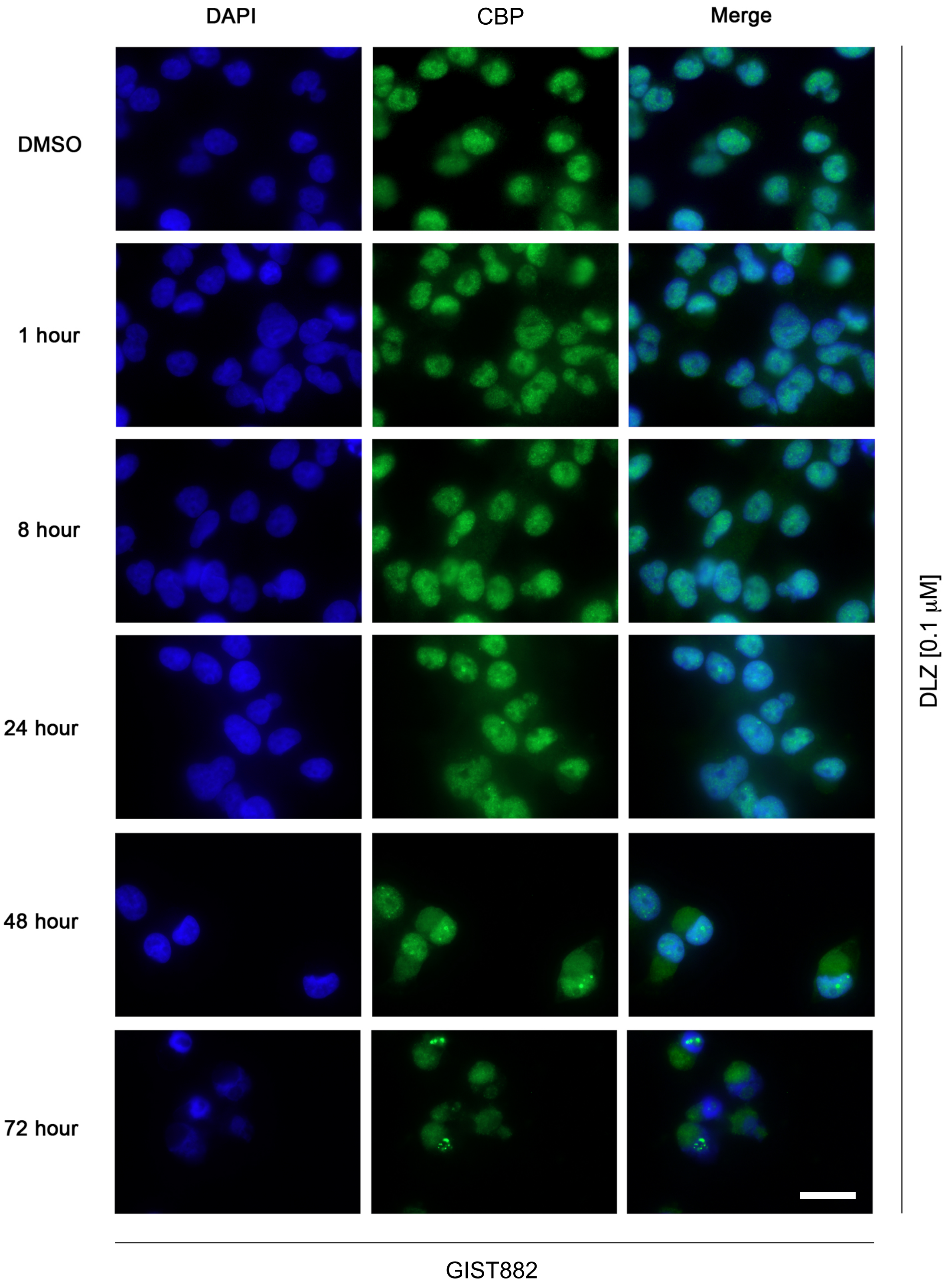


Rausch *et al.*, Supplemental Figure 7

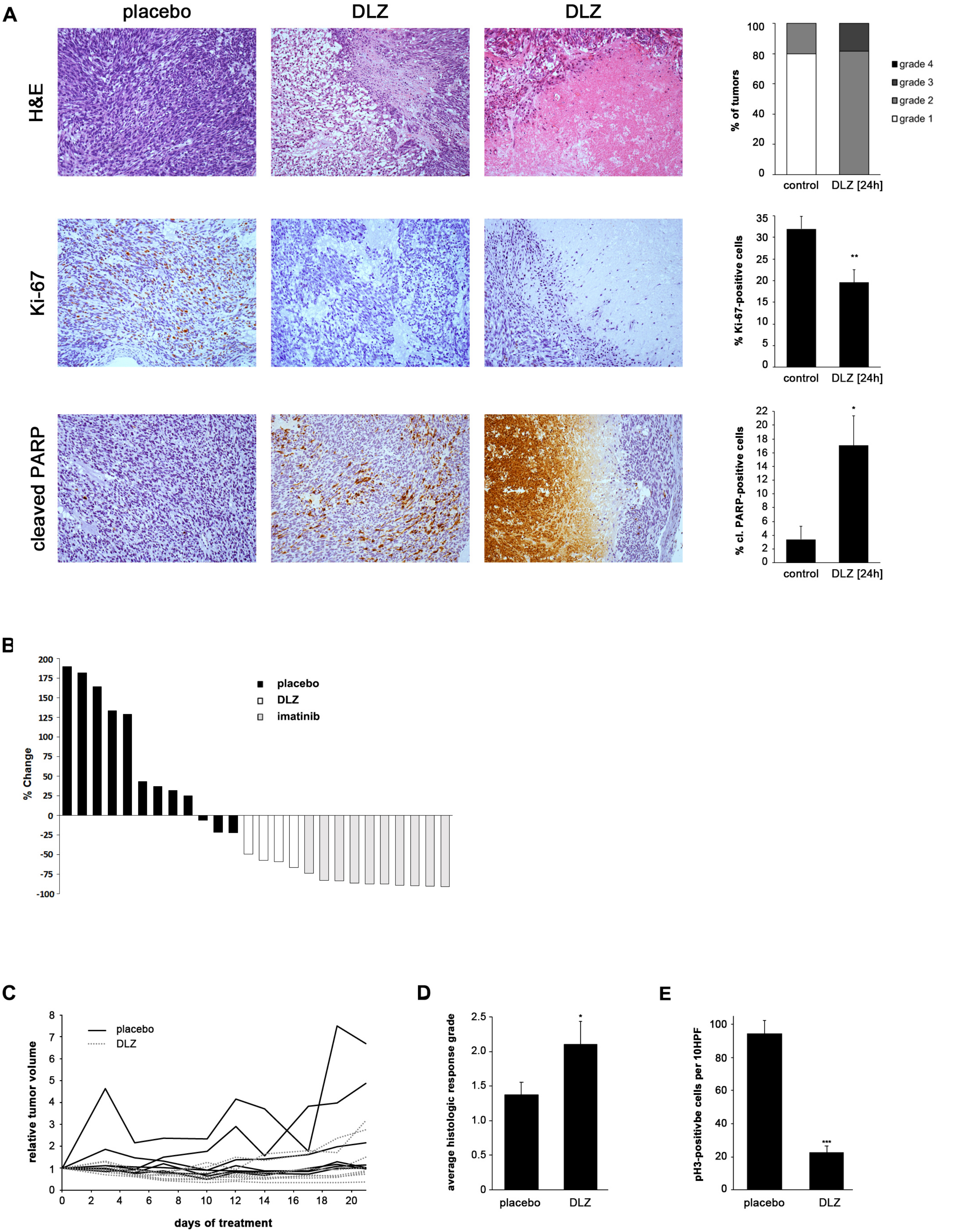


Rausch *et al.*, Supplemental Figure 8
